# Image-based tracking of ripening in wheat cultivar mixtures: a quantifying approach parallel to the conventional phenology

**DOI:** 10.1101/239798

**Authors:** Abbas Haghshenas, Yahya Emam

## Abstract

The lack of quantitative methods independent of the conventional qualitative phenology, may be a vital limiting factor to evaluate the temporal trends in the crop growth cycle, particularly in the heterogeneous canopies of cultivar mixtures. A digital camera used to take ground-based nadir images during two years of a field experiment conducted at the College of Agriculture, Shiraz University, Iran; in 2014-15 and 2015-16. The experimental treatments consisted of 4 early- to middle-ripening wheat cultivars and their 10 mixtures, under post-anthesis well- and deficit-irrigation conditions, arranged in a randomized complete block design with 3 replicates. Then the images were processed and three image-derived indices including CC (canopy cover), GR [(G-R/G); RGB color system], and CCGR (CC×GR) were used as the quantifying criteria. The declining trends of these indices during ripening showed strong fits to binomial equations, based on which simple prediction models were suggested and validated. Furthermore, the split linear trends and their slopes were estimated to assess the short-term variations. Some agronomic aspects were also evidenced using the mixtures-monoculture diversions, and the relationship between CC and GR. The frameworks evaluated appears to provide the reliable and simple solutions for quantifying the crop temporal trends parallel to the conventional phenology.

## Introduction

Monitoring or predicting the irreversible trend of successive events in crop growth and development (i.e. phenology), is a fundamental necessity on almost every field crop study or practice, even where it is not the main objective. Conventional methods in phenological studies-e.g. Feekes (Feekes, 1941; Large, 1954), Zadoks (Zadoks *et al*., 1974), and BBCH scales (Lancashire *et al*., 1991)-are mainly based on professional qualitative descriptions; even though they use numerical bases (Landes and Porter, 1989), and/or reporting the crop stages based on some quantitative aspects such as 50% flowering (which usually depends on the observer’s perception of the canopy status, instead of being the result of an exact counting of the plants). However, especially where the high degree of accuracy in estimations are needed, e.g. in developing or running crop models, an error of one or two day(s) in distinguishing the phenological events /or periods may lead to considerable miscalculations (e.g. in calculating the base temperature and growth degree days –GDD-using current formulae-Yang *et al*., 1995-, particularly when the data is limited to few plantings). Obviously, the difficulties and limitations in this context are associated with two main facts: (a) the continuous trend of growing and developmental processes in single plants, which are characterized in the form of discrete qualitative codes; and (b) the phenological differences between individual plants within a canopy, even in the most homogenous stands of monocultures consisted of the modern pure genotypes.

Cultivar mixtures are investigated during the recent decades as the potential alternatives for conventional intense cropping systems (Kiær *et al*., 2009; Borg *et al*., 2017; Reiss and Drinkwater, 2017), mainly due to the expected advantages of improving biodiversity, based on the ecological principles. If the cultivar mixtures are designed based on the phenological differences of the included components (i.e. cultivars), determination of the crop phenology in such a heterogeneous population-as a whole canopy-would even be a more challenging problem, so that the efficient determination of the crop stage by the conventional methods appears to be impossible. For instance, Haghshenas *et al*. (2013) and Fang *et al*. (2014) evaluated the wheat cultivar mixtures with different ripening patterns aiming to increase water use efficiency under water-limited conditions. In these situations, even after distinguishing the cultivars in the mixtures, the way of reporting the overall crop phenology will be problematic, in the absence of an appropriate estimation framework.

Remote sensing approaches are currently the well-established tools for monitoring and predicting crop status in large- to farm-scales. Accordingly, there is a considerable number of studies in the literature reporting results of crop phenology recognition or modeling using remote sensing techniques, mostly based on the well-known spectral indices (e.g. NDVI, Normalized Difference Vegetation Index) and sensors (Sakamoto *et al*., 2010; Lopes and Reynolds, 2012; Lausch, *et al*., 2015; Aubrecht *et al*., 2016; Magney *et al*., 2016; Zeng *et al*., 2016; Canisius *et al*., 2017; Gao *et al*., 2017; Liu *et al*., 2017). Among the reliable, readily available, and low cost sensors, are common commercial digital cameras, which are increasingly attracting attentions in crop sciences by providing robust relationships between image-derived indices and bio-physiological criteria (Li *et al*., 2010; Sakamoto *et al*., 2012; Wang *et al*., 2013; Lee and Lee, 2013; Hunt Jr *et al*., 2013; Easlon and Bloom, 2014; Zou *et al*., 2014). Despite the novel multi-to hyperspectral sensors and criteria developed, digital cameras seem to be capable to remain as a desirable choice for determining the crop status quantitatively and accurately, due to having high spatial and color resolutions, and considerable overlap between the spectral ranges of visible light and photosynthetically active radiation (PAR, McCree, 1972). However, despite the relatively more frequent reports for forest and rangeland species (Bradley *et al*., 2010; Ide and Oguma, 2010; Granados *et al*., 2013; Henneken *et al*., 2013; Alberton *et al*., 2014; Inoue *et al*., 2014; Alberton *et al*., 2017; Lang *et al*., 2017; Toda and Richardson, 2017), studies with the purpose of using digital color images for evaluating the crop phenology are rare (Sakamoto *et al*., 2011; Imukova *et al*., 2015; Bargiel, 2017).

Furthermore, even in the remote sensing approaches, the quantitative outputs are usually reported based on the conventional qualitative phenological events, and there are few reports in the literature in which the amounts of a quantitative index were taken independently as the crop developmental “events”, themselves (Lopes and Reynolds, 2012). In the study of Lopez and Reynolds (2012), expression of stay-green was evaluated for a considerably diverse set of wheat populations, based on NDVI values at physiological maturity. They suggested the rate of senescence regressed on degree days, as an independent measurement of stay-green without the confounding effect of phenology. Such attempts may be considered as the evidences for the inevitable necessity of developing or generalizing novel approaches in order to fill the gap between the currently available qualitative scales, and the increasing need for determining the phenological events more precisely, efficiently, and quantitatively.

Therefore, in an ideal horizon, the crop temporal dynamics caused by various phenomena with distinct biological bases e.g. tillering, flowering, or ripening would be alternatively represented using a unique mathematical terminology. For instance, if supported by robust evidences and shown by adequate studies, a researcher would report that a given wheat cultivar needs “n” GDDs to reach its maximum amount of the CC (the image-derived canopy cover index) under optimal conditions, without necessarily referring to its conventional qualitative phenology (e.g. reporting the growth stage was at the middle anthesis, or the Zadoks code 65). Obviously, achieving this goal requires that the remote sensing-based indices (i) reflect the crop variations over the time (or against thermal time/ or GDD) appropriately, and (ii) be predictable enough to be used in the crop models, as the alternatives to the conventional phenological events.

The objectives of the present study were: (1) monitoring and quantifying the ripening trends in monocultures and mixtures of four winter wheat cultivars with different ripening patterns under well-and deficit-irrigation conditions, utilizing image-derived indices; and (2) evaluating the option of developing simple independent quantitative frameworks parallel to the conventional qualitative ones for identifying and modeling the crop trends over the season, employing uncomplicated image-based computable criteria.

## Materials and methods

In order to evaluate the ripening trends in various mono- and mixed cropping of 4 early to middle ripening winter wheat cultivars, series of digital images were taken during two growing seasons and processed.

### Field experiments

A 2-year factorial field experiment was conducted during 2014–15 and 2015–16 growing seasons at the research field of College of Agriculture, Shiraz University, Iran (29°73′ N latitude and 52°59′ E longitude at an altitude of 1,810 masl). Treatment were included the 15 mixing ratios of four early to middle ripening wheat cultivars [Chamran (1), Sirvan (2), Pishtaz (3), and Shiraz (4), respectively] including the 4 monocultures and their every 10 possible mixtures, which were grown with 3 replicates under two normal and post-anthesis deficit-irrigation conditions. The experimental design was RCBD (Randomized Complete Block Design) in which all the 90 (2×2 meter) plots were arranged in a lattice configuration with 1 meter distances. Plant density was set to 450 plants/m^2^ and seeds were mixed with equal ratios (1:1-1:1:1- and 1:1:1:1, for the 2-, 3-, and 4-component blends, respectively) considering their 1000-grain weights and germination percentages. The planting dates in the first and second growing seasons were November 20 and November 5, respectively; and based on the soil test, only 150 kg nitrogen/ha (as urea) was applied in three equal splits i.e. at planting, early tillering, and anthesis. No pesticide was used and weeding was done by hand.

Based on the local practices, irrigation interval was set at ten days, and the amount of irrigation water was estimated using Fao-56 Penman-Monteith model with local corrected coefficients (Razzaghi and Sepaskhah, 2012; Shahrokhnia and Sepaskhah, 2013) which was reduced to 50% of evapo-transpirational demand from the first irrigation after anthesis.

### Imaging

The nadir images of plots were taken throughout the both growing seasons in the same way i.e. from 150 cm above the soil surface during the period between solar noon and 2 hours later. The imaging events were more frequent from flowering towards the end of season, due to more rapid changes in the canopy status. Images were taken by a common commercial digital camera (Canon PowerShot SX100 *IS*), setting to auto mode and the maximum imaging resolution of 8.0 megapixels. The overall imaging duration for each day was maximum 40 minutes.

### Image processing and indices

Image processing was carried out using an exclusive MATLAB code, by which the images were primarily segmented into two *“green vegetation”* and background parts based on the common thresholding formula of G-R>0 (Wang *et al*., 2016; G and R stand for green and red color values in RGB color system, respectively). Subsequently, the following image-derived indices were calculated for each image:

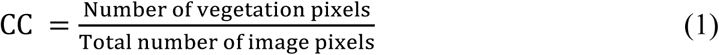

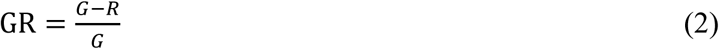

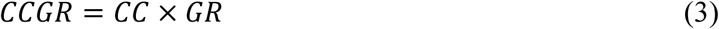
where *CC* is canopy cover (Guevara-Escobar *et al*., 2005; Wang *et al*., 2016), and “*G*” and “*R*’ are mean green and red values of vegetation parts (pixels) in RGB color system, respectively, with the range of 0–255 (i.e. for zero to maximum possible reflection recorded for each color). It is notable that CC is a well-known criterion related to the quantitative development of the green canopy; while Gft is the normalized amount of G-R (Wang *et al*., 2013), a quantitative measure associated with quality of the canopy spectral behavior, regardless of its size. Indeed, the GR index shows the difference between the recorded red and green values in each vegetation image pixel, independent of the size of canopy coverage (i.e. the comparative quality of light per area unit of canopy cover). Therefore, the GR index may be taken as an indicator for quality of the photosynthesis apparatus; as is expected, the difference between red and green reflection from the canopy would be larger in more desirable and healthy conditions. Furthermore, the CCGR index may provide an overall integrative estimation of the quantity and quality of reflection from green surfaces with respect to the image area (i.e. per occupied ground surface). Obviously, each of the three mentioned indices have a theoretical range between 0 to 1. The trend of variations in indices were evaluated during the season (particularly from anthesis) using simple linear or various binomial equations.

The diurnal temperatures for calculating accumulated thermal time (ATT) and growth degree days (GDD) were obtained from the weather station located about 500 meters from the experimental field. The Thermal time was calculated by summing the average diurnal temperatures (°C) in the certain period (in the most cases, from sowing to ripening i.e. CC=0); and the individual base temperature for each cultivar was estimated based on the “*standard deviation in days, SD*” equation (Yang *et al*., 1995) using the data recorded for the two growing seasons (whose results were also exactly as the same as the “*coefficient variation, CV*” and the “*regression coefficient, RE*” methods, due to using the data of two plantings in the equations). To the best of our knowledge, it is the first time that the base temperature is calculated on the basis of an image-derived date (event) i.e. when the CC of each cultivar was reached zero, as the results of binomial equations between the DAS (days after sowing) and CC values, where they were the independent and the dependent variables, respectively. Thereafter, GDD of each cultivar was calculated for the defined periods (sowing to ripening i.e. DAS0 to DAScc=0), by the following equation (McMaster and Wilhelm, 1997):

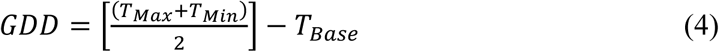

where if[(*T_Max_ + T_Min_*)/2] < *T_Base_*, then [(*T_Max_ + T_Min_*)/2] = *T_Base_*.

Model validations were performed by testing the data of the second year as the independent inputs of the equations obtained from the first year, and consequently comparing the RMSE (root-mean-square error).

The mathematical analyses (equation fitness) were carried out using Origin Pro 8 software (OriginLab, Northampton, MA) and XLSTAT Version 2016.02.28451 (Addinsoft). Statistical analyses were carried out using IBM SPSS Statistics for Windows, Version 19.0 (Armonk, NY: IBM Corp.) and mean comparisons were performed using LSD and Tukey’s tests. Finally, all charts and figures were made and edited by Microsoft Excel 2016 and Adobe Photoshop CC 2017.

## Results

The comparative differences in the ripening trends of the four monocultures under the well-irrigation conditions of the 1^st^ year are shown in Fig. 1, using the original images, numerical quantities, and also in a novel kind of diagram. As the values of CC, GR, and CCGR indicate, these criteria have declined towards the end of the season; the trends whose rates also decrease from the monoculture of the early-ripened cultivar to the middle-ripening one. The effects of irrigation and mixture treatments on the three image-derived criteria were also significant in most of the imaging dates in both years, particularly from middle to late season (Table S1). In general, post-anthesis deficit-irrigation reduced the CC, GR, and CCGR values. Differences among the mixtures were smaller under deficit-irrigation, compared with the well-irrigation conditions (Table S2 & S3). The theoretical range of either criterion is between 0 to 1 (M&M); however, the actual records were as below: the highest amounts of CC recorded in the 1^st^ and 2^nd^ years were 0.858 (175 DAS) and 0.933 (194 DAS), respectively; similar records for GR were 0.259 (155 DAS) and 0.237 (174 & 191 DAS); and were equal to 0.218 (155 DAS) and 0.221 (191 DAS) for CCGR. Therefore, the actual (observed) ranges of GR and CCGR were between 0 to the maximum amount of 0.3.

**Figure 1.**
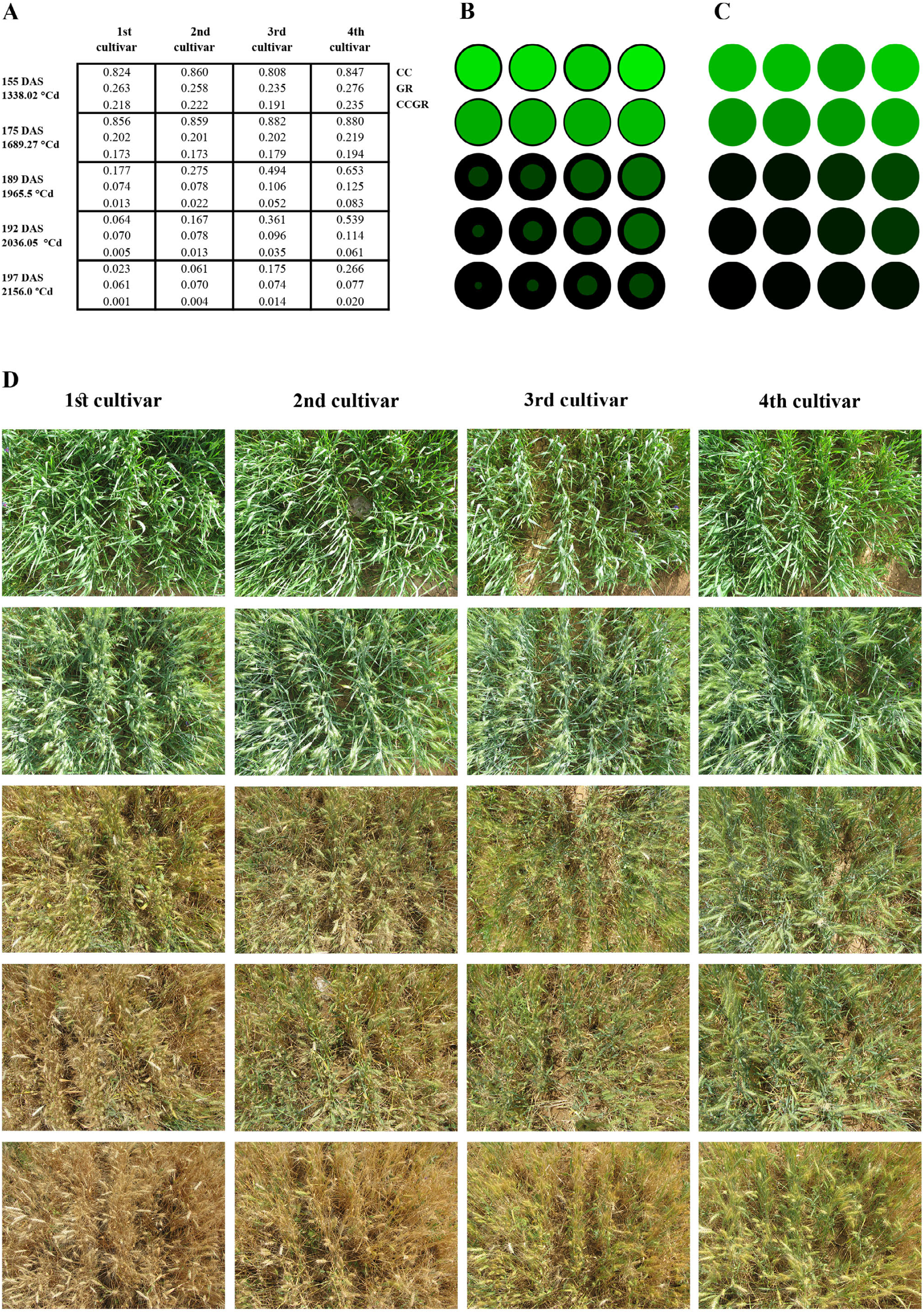
The ripening trends in the 4 monocultures of early to middle ripening wheat cultivars during the 1^st^ season, quantified using the CC, GR, and CCGR image-derived criteria. (A) The table represents the labels and the quantities of the image-derived criteria late in the season. The time of imaging events are shown on the basis of DAS (days after sowing) and thermal time, at the left side of the table. The configuration of the images and objects in other parts of the figure also follow this table. (B) The visualized concepts of CC and GR criteria. The equal-sized black circles in the background represent the unit of area (in the image and/or on the ground); the comparative size and color ratios of 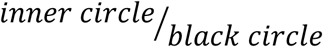 indicate the CC and GR concepts, respectively, which are drawn based on the real ratios; so the inner circles with higher degrees of greenness (i.e. also look brighter) represents the images with higher GRs. The RGB color for drawing inner circles are determined as: Red=0, Blue=0, Green= GR × a constant value (i.e. 850; ≅255×3.333; for strengthening and making the color more visible); thus, the actual ratios are kept constant. (C) The visualized concept of CCGR, which implies distributing the overall green content of each inner circle across the black circle in the part B (or diluting the normalized greenness of the vegetation parts based on the ratio of canopy cover). Furthermore, the CCGR seems to be recognizable in the part B, as the overall perception of brightness vs. darkness of the circle pairs. (D) The reduced-size original images of the experimental plots whose calculated criteria are represented in the table part A, and simulated in the parts B and C.

### The binomial declining trends during ripening

Evaluating the declining trend of each criterion based on thermal time during ripening, revealed that the simple form of binomial models had strong fits to the data (Fig. 2). Among the image-derived indices, the GR trend had a relatively more gradual declining slope compared with CC and CCGR, irrespective the irrigation condition and season; so that based on the binomial equation estimations, the terminal GR values became zero (GR_0_) later than CC, i.e. it needed more accumulated thermal time (Fig. 2 and Fig. S1). In other words, the term “GR_0_” remains as a theoretical concept, because CC has become zero formerly and there are not green points (pixels) in the canopy any longer, whose spectral quality might be evaluated using GR. Accordingly, in this study, CC_0_ (CC=0) was selected as the ripening date or terminal point, based on the objectives (CCGR_0_ also may be alternatively chosen as the canopy terminal date, where the objectives require). Although CCGR has comparatively smaller values than the corresponding CC and GR (because is the product of them, both of which have values less than 1), its zero values (CCGR_0_) at the late season estimated by the binomial trends generally need ATTs around or more than that is required for CC_0_, with comparatively less differences between cultivars (Fig. 2).

**Figure 2.**
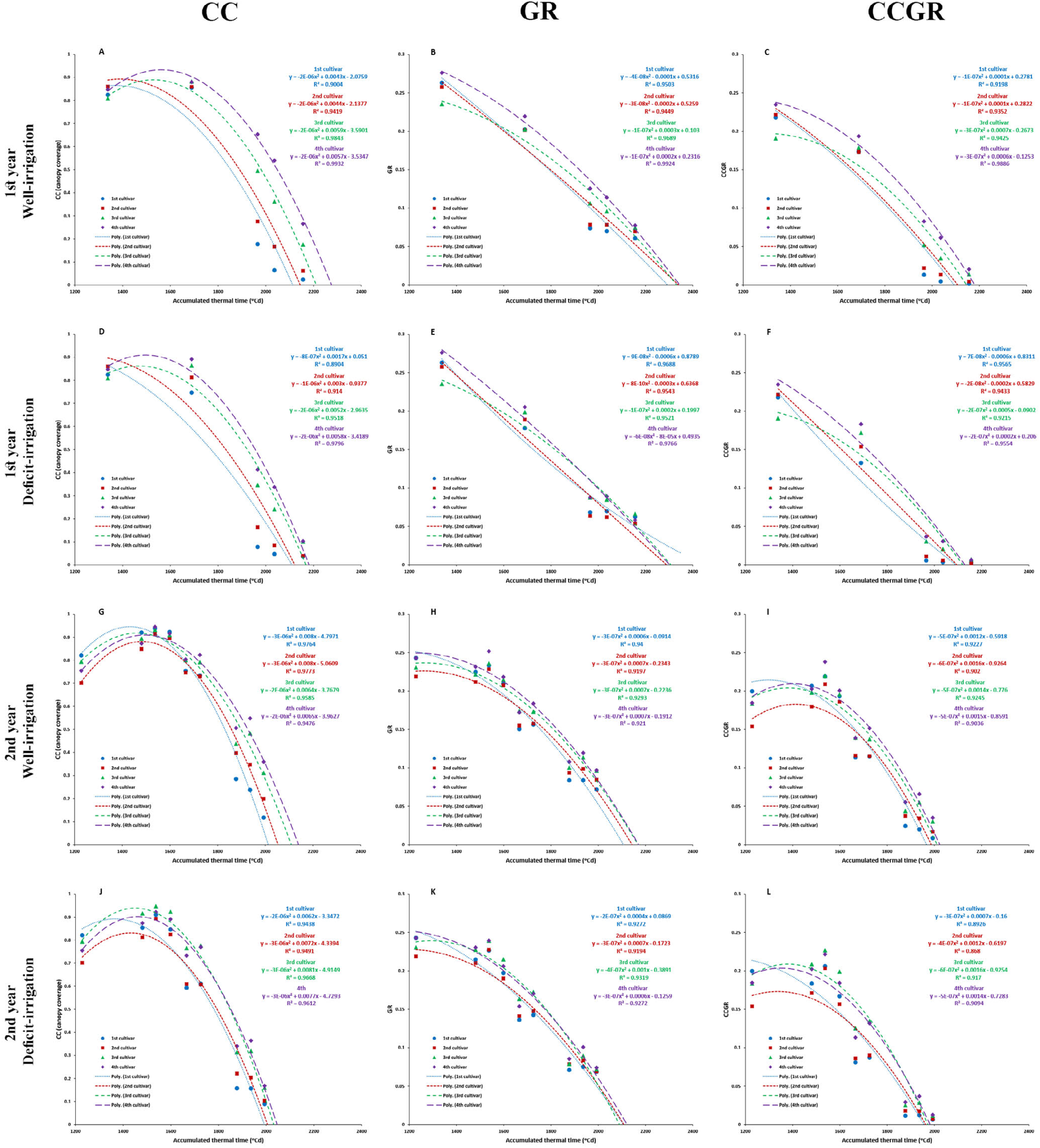
Comparative trends of the image-derived indices under well- and deficit-irrigation conditions, during the ripening period of the firâ and second years.

The predicted thermal times of ripening for the 4 cultivars (CC_0_) estimated based on the binomial equations showed comparatively broader ranges under well-irrigation condition in both years, while they seem to be more similar (had a narrower range), under the deficit-irrigation conditions (Fig. 2). Such trend is recognizable in Fig. 3 which represents the comparative declining trends of CC for every 15 mixture treatments, under different irrigation conditions over the two years. As a general rule –irrespective of season or condition-, the most early- and late-ripening cultivars (i.e. the 1^st^ and 4^th^ cultivars) had the fastest and slowest binomial declining trends during ripening, respectively, and thus eventually had the minimum and maximum ATT extrema at ripening (CC_0_); the range in which the other monocultures and mixtures were placed. Again, the effect of post-anthesis deficit-irrigation is obvious in the form of decreasing diversities among the ripening events (i.e. the smallest range) of the mixture treatments (Fig. 3). Moreover, the deficit-irrigation significantly accelerated the ripening rate, and consequently reduced the ATTs needed for the CC_0_ event (Fig. 3, Tables S4 to S6). The effects of mixtures and irrigation treatments on the ripening trends, ATTs required for reaching the maximum CC, and for CC_0_ were significant (Table S4); such that the differences between ripening of the early- to middle-ripening cultivars -and also among the mixtures-were significant. Furthermore, the R^2^ and RMSE values of the binomial trends were affected significantly by the irrigation treatment, that generally reduced the regression fit under deficit-irrigation conditions.

**Figure 3.**
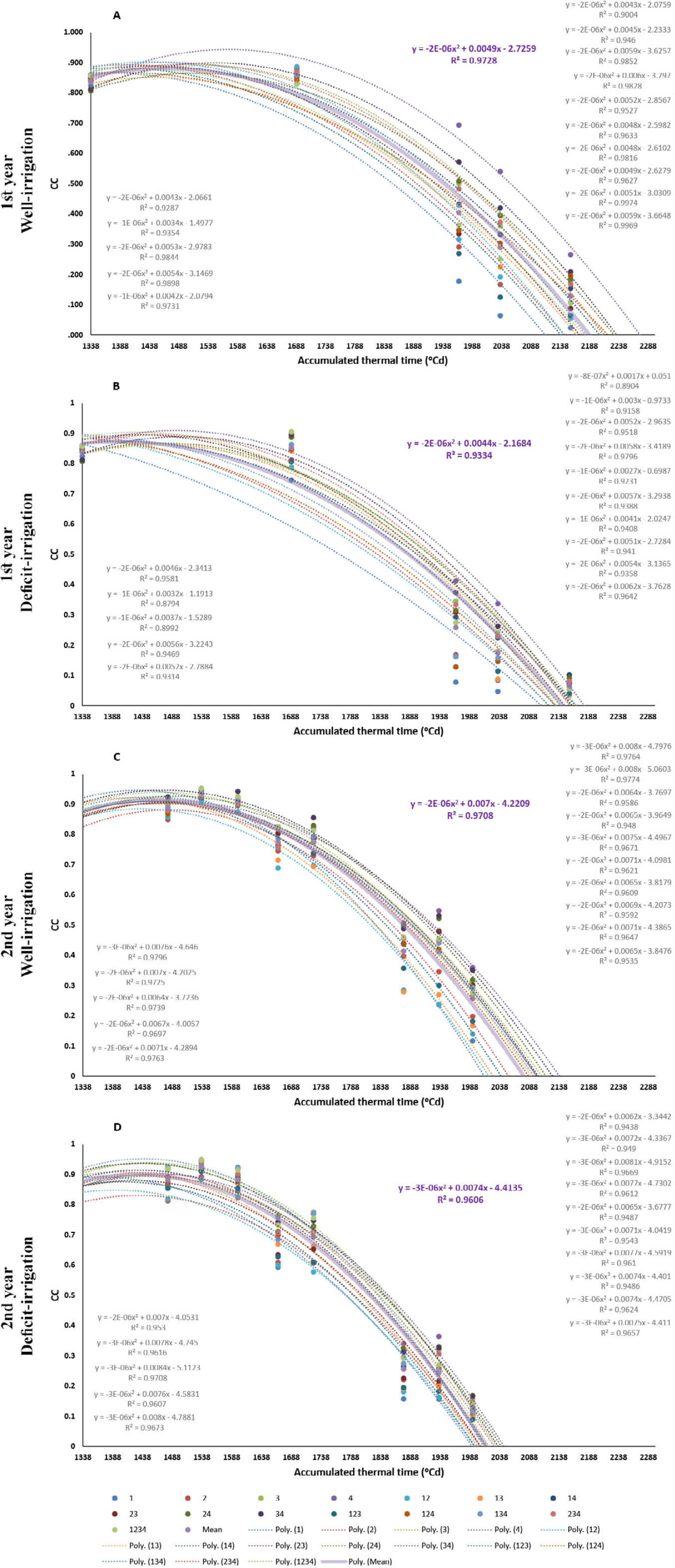
The binomial declining trends of CC (canopy cover) during the ripening of the mixture treatments of 4 wheat cultivars, under well- and post-anthesis deficit-irrigation conditions. Mixture treatments include the monocultures and mixtures of the four early to middle ripening wheat cultivars. Accordingly, each digit stands for a single cultivar included in the mixture. **A** and **B** show the trends in the first year; and **C** and **D** indicate the 2nd year’s trends. The thick purple curve and the corresponding bold equation, show the average trend over the treatments, and the equations arranged from top to down, represent the trends of the treatments in the order of below: 1,2, 3, 4, 12, 13, 14, 23, 24, 34, 123, 124, 134, 234, 1234; where 1, 2, 3, and 4 are the monocultures of the early to middle ripening cultivars, respectively, and the other treatments are the mixtures included these cultivars, e.g. the treatment 1234 is the 4-component mixture of the 4 cultivars.

In order to evaluate the option of using the simple binomial models for predicting the amounts of the image-derived indices based on thermal time, and particularly estimating the time of the terminal CC_0_, GR_0_, and CCGR_0_ events, the equations of the first year were used for predicting the 2^nd^ year’s dataset. Tables 1 to 3, along with the Fig. S2 to S7 represent the results of the model validations based on accumulated thermal time, and the Fig. S8 show the outcomes of the estimations based on cultivar growing degree days. Among the three indices, CC had the best model fitness (R^2^ and RMSE), and the least deviation in prediction of the date or ATT of the terminal zero point (among CC_0_, GR_0_, and CCGR_0_); however, the deviation of its regression line from the 1:1 line was more than GR (see the intercepts and slopes in the Tables 1 to 3, and also compare the corresponding trends in Fig. S2 to S7). Moreover, it appeared that almost all the regression parameters were influenced by the deficit-irrigation negatively, irrespective the type of the image-derived index. For instance, the average deviations in the predicted CC_0_, GR_0_, and CCGR_0_ events showed a gradual raise from 5.2 to 6.2, 9.3 to 10.2, and 6.5 to 6.9 days, respectively, due to the post-anthesis deficit-irrigation.

Figure S8 represents the results of the model validation for the binomial models in which the image-derived indices are regressed against GDD, instead of ATT. As mentioned before, the base temperatures and GDDs were calculated based on the sowing-to-CC_0_ periods in both seasons. The calculated growth degree days for the well-irrigated monocultures were 3001.8, 3107.3, 3249, 3733.3 for the 1^st^, 2^nd^, 3^rd^, and the 4^th^ cultivars, using the base temperatures equal to -4.6, -4.8, -5.1, and -7.1 °C, respectively (calculations are not shown).

Obviously, in this method the diversions from the 1:1 line are reduced compared with utilizing ATT (see the slopes and intercepts in the Fig. S8). However, since the calculations of the base temperatures were limited to the two plantings of the present study, the results should be interpreted and generalized with caution.

### The Linear trends

Although the best fitting regression trends for the image-derived indices against thermal time were binomial, the linear trends may also provide valuable information particularly for the short-term variations. Figure 4 shows the split linear trends of variations in the image-derived indices of the 1^st^ cultivar monoculture. Based on the objectives, the equations of the linear trends between pairs of points (observations) may be used for interpreting the variation through the season. For instance, despite the fact that the number of imaging events in the two seasons were not the same, and also they were not necessarily synchronized, the linear increasing trends from sowing to the maximum CC observed (CC_max_) were approximately similar; with the slopes equal to 0.0005 vs 0.0006 (Fig. 4, see the dotted lines in the parts A and B). The CC linear declining trends of this early-ripening cultivar were still more identical; having the slopes exactly equal to -0.0018, and the intercepts of 3.87 vs 3.71 in the first and second year, respectively. It implies that the declining rates of CC during ripening was in average, about 3 times faster than their increasing rate during the canopy development. The linear GR declining trends were also very similar in the both growing seasons, though the linear increasing trend of the first year was 2.5 times faster, compared with the second year. A relatively comprehensive comparisons were carried out for other monocultures and mixtures, in the same way (Fig. S9, Tables S7 to S9).

**Table 1.** Validation of the binomial model for prediction of the CC (canopy cover) trends against thermal time, during ripening of cultivar mixtures under well- and deficit-irrigation conditions.

| Moisture condition | Mixture treatments | $R^2$ | RMSE | Model P-value | Intercept | Slope | Pred.-Obs. thermal time to $CC_0$ ($^{\circ}Cd$) | Pred. -Obs. days to $CC_0$ (day) |
| --- | --- | --- | --- | --- | --- | --- | --- | --- |
| Well irrigation | 1 | 0.964 | 0.066 | < 0.0001 | -0.289 | 1.447 | 100.39 | 4.8 |
|  | 2 | 0.938 | 0.070 | < 0.0001 | -0.198 | 1.219 | 91.98 | 4.4 |
|  | 3 | 0.897 | 0.080 | 0.000105 | -0.381 | 1.447 | 102.42 | 4.9 |
|  | 4 | 0.812 | 0.095 | 0.000912 | -0.445 | 1.422 | 135.35 | 6.5 |
|  | 12 | 0.967 | 0.058 | < 0.0001 | -0.424 | 1.449 | 125.09 | 6.0 |
|  | 13 | 0.958 | 0.067 | < 0.0001 | -0.507 | 1.670 | 139.00 | 6.7 |
|  | 14 | 0.960 | 0.050 | < 0.0001 | -0.333 | 1.408 | 104.59 | 5.0 |
|  | 23 | 0.950 | 0.055 | < 0.0001 | -0.163 | 1.205 | 68.10 | 3.3 |
|  | 24 | 0.957 | 0.050 | < 0.0001 | -0.405 | 1.538 | 114.33 | 5.5 |
|  | 34 | 0.866 | 0.088 | 0.000274 | -0.399 | 1.483 | 107.62 | 5.2 |
|  | 123 | 0.960 | 0.061 | < 0.0001 | -0.246 | 1.324 | 98.42 | 4.7 |
|  | 124 | 0.938 | 0.064 | < 0.0001 | -0.285 | 1.382 | 132.69 | 6.4 |
|  | 134 | 0.971 | 0.044 | < 0.0001 | -0.273 | 1.325 | 81.76 | 3.9 |
|  | 234 | 0.953 | 0.054 | < 0.0001 | -0.423 | 1.477 | 118.07 | 5.7 |
|  | 1234 | 0.958 | 0.055 | < 0.0001 | -0.199 | 1.301 | 87.44 | 4.2 |
|  | Mean | 0.937 | 0.064 | - | -0.331 | 1.406 | 107.15 | 5.2 |
| Deficit irrigation | 1 | 0.895 | 0.117 | 0.000114 | -0.187 | 1.317 | 117.32 | 5.6 |
|  | 2 | 0.896 | 0.103 | 0.000109 | -0.213 | 1.213 | 116.26 | 5.6 |
|  | 3 | 0.964 | 0.062 | < 0.0001 | -0.521 | 1.690 | 136.27 | 6.6 |
|  | 4 | 0.950 | 0.067 | < 0.0001 | -0.514 | 1.552 | 135.51 | 6.5 |
|  | 12 | 0.914 | 0.099 | < 0.0001 | -0.251 | 1.284 | 125.54 | 6.0 |
|  | 13 | 0.934 | 0.091 | < 0.0001 | -0.515 | 1.625 | 132.80 | 6.4 |
|  | 14 | 0.957 | 0.070 | < 0.0001 | -0.481 | 1.606 | 150.03 | 7.2 |
|  | 23 | 0.948 | 0.077 | < 0.0001 | -0.493 | 1.531 | 141.49 | 6.8 |
|  | 24 | 0.958 | 0.062 | < 0.0001 | -0.423 | 1.513 | 118.21 | 5.7 |
|  | 34 | 0.926 | 0.088 | < 0.0001 | -0.506 | 1.607 | 125.21 | 6.0 |
|  | 123 | 0.953 | 0.075 | < 0.0001 | -0.483 | 1.553 | 144.43 | 7.0 |
|  | 124 | 0.914 | 0.098 | < 0.0001 | -0.250 | 1.345 | 118.74 | 5.7 |
|  | 134 | 0.937 | 0.089 | < 0.0001 | -0.314 | 1.451 | 122.18 | 5.9 |
|  | 234 | 0.955 | 0.068 | < 0.0001 | -0.475 | 1.537 | 127.87 | 6.2 |
|  | 1234 | 0.967 | 0.062 | < 0.0001 | -0.417 | 1.499 | 119.85 | 5.8 |
|  | Mean | 0.938 | 0.082 | - | -0.403 | 1.488 | 128.78 | 6.2 |

Mixture treatments include the monocultures and mixtures of the four early- to middle-ripening wheat cultivars. Accordingly, each digit stands for a single cultivar included in the mixture, so the treatment 1234 shows the 4-component mixtures of cultivars 1,2,3 and 4. $R^2$ , $RMSE$ , $Model\ P\text{-}value$ , $Intercept$ , and $Slope$ are the parameters of the linear regression between the predicted and observed values. $CC_0$ is the time (and/or thermal time) when CC becomes zero (at ripening). " $Pred.\text{-}Obs.$ " shows the difference between the predicted and observed values. Number of days in the last column are estimated by dividing the corresponding thermal times (in the adjacent column) by the averaged diurnal thermal times of the final 10 days.

**Table 2.** Validation of the binomial model for prediction of the GR trends against thermal time, during ripening of cultivar mixtures under well- and deficit-irrigation conditions.

| Moisture condition | Mixture treatments | $R^2$ | RMSE | Model P-value | Intercept | Slope | Pred.-Obs. thermal time to $GR_0$ ($^{\circ}Cd$) | Pred. -Obs. days to $GR_0$ (day) |
| --- | --- | --- | --- | --- | --- | --- | --- | --- |
| Well irrigation | 1 | 0.899 | 0.024 | 0.000100 | -0.020 | 1.004 | 186.70 | 9.0 |
|  | 2 | 0.844 | 0.025 | 0.000468 | 0.003 | 0.874 | 195.31 | 9.4 |
|  | 3 | 0.914 | 0.018 | < 0.0001 | -0.048 | 1.209 | 164.04 | 7.9 |
|  | 4 | 0.900 | 0.020 | < 0.0001 | -0.041 | 1.060 | 172.36 | 8.3 |
|  | 12 | 0.890 | 0.023 | 0.000133 | -0.020 | 0.946 | 194.89 | 9.4 |
|  | 13 | 0.905 | 0.022 | < 0.0001 | -0.054 | 1.186 | 225.41 | 10.9 |
|  | 14 | 0.901 | 0.021 | < 0.0001 | -0.023 | 1.021 | 198.64 | 9.6 |
|  | 23 | 0.862 | 0.021 | 0.000303 | 0.008 | 0.890 | 162.98 | 7.8 |
|  | 24 | 0.879 | 0.023 | 0.000188 | -0.023 | 1.056 | 213.53 | 10.3 |
|  | 34 | 0.903 | 0.021 | < 0.0001 | -0.052 | 1.206 | 197.21 | 9.5 |
|  | 123 | 0.866 | 0.024 | 0.000268 | 0.004 | 0.924 | 176.93 | 8.5 |
|  | 124 | 0.882 | 0.022 | 0.000174 | -0.008 | 0.950 | 269.76 | 13.0 |
|  | 134 | 0.922 | 0.019 | < 0.0001 | -0.021 | 1.017 | 169.27 | 8.1 |
|  | 234 | 0.923 | 0.018 | < 0.0001 | -0.039 | 1.080 | 175.35 | 8.4 |
|  | 1234 | 0.869 | 0.024 | 0.000250 | -0.006 | 0.974 | 219.42 | 10.6 |
|  | Mean | 0.891 | 0.022 | - | -0.023 | 1.026 | 194.787 | 9.376 |
| Deficit irrigation | 1 | 0.867 | 0.027 | 0.000266 | -0.005 | 0.935 | 344.55 | 16.6 |
|  | 2 | 0.847 | 0.026 | 0.000432 | 0.005 | 0.861 | 181.25 | 8.7 |
|  | 3 | 0.894 | 0.024 | 0.000119 | -0.066 | 1.309 | 217.44 | 10.5 |
|  | 4 | 0.887 | 0.024 | 0.000149 | -0.019 | 0.968 | 180.24 | 8.7 |
|  | 12 | 0.877 | 0.025 | 0.000200 | -0.005 | 0.882 | 221.73 | 10.7 |
|  | 13 | 0.920 | 0.021 | < 0.0001 | -0.071 | 1.255 | 194.54 | 9.4 |
|  | 14 | 0.877 | 0.028 | 0.000202 | -0.043 | 1.098 | 248.82 | 12.0 |
|  | 23 | 0.883 | 0.024 | 0.000167 | -0.037 | 1.053 | 193.69 | 9.3 |
|  | 24 | 0.876 | 0.025 | 0.000204 | -0.019 | 1.025 | 177.59 | 8.5 |
|  | 34 | 0.915 | 0.023 | < 0.0001 | -0.068 | 1.252 | 190.39 | 9.2 |
|  | 123 | 0.912 | 0.022 | < 0.0001 | -0.069 | 1.174 | 226.92 | 10.9 |
|  | 124 | 0.861 | 0.028 | 0.000310 | -0.010 | 0.967 | 218.20 | 10.5 |
|  | 134 | 0.872 | 0.029 | 0.000231 | -0.026 | 1.071 | 230.86 | 11.1 |
|  | 234 | 0.897 | 0.023 | 0.000107 | -0.030 | 1.049 | 176.04 | 8.5 |
|  | 1234 | 0.889 | 0.027 | 0.000137 | -0.044 | 1.114 | 176.47 | 8.5 |
|  | Mean | 0.885 | 0.025 | - | -0.034 | 1.067 | 211.91 | 10.2 |

Mixture treatments include the monocultures and mixtures of the four early- to middle-ripening wheat cultivars. Accordingly, each digit stands for a single cultivar included in the mixture, so the treatment 1234 shows the 4-component mixtures of cultivars 1,2,3 and 4. $R^2$ , $RMSE$ , $Model\ P\text{-}value$ , $Intercept$ , and $Slope$ are the parameters of the linear regression between the predicted and observed values. $GR_0$ is the time (and/or thermal time) when GR becomes zero (at ripening). " $Pred.\text{-}Obs.$ " shows the difference between the predicted and observed values. Number of days in the last column are estimated by dividing the corresponding thermal times (in the adjacent column) by the averaged diurnal thermal times of the final 10 days.

**Table 3.** Validation of the binomial model for prediction of the CCGR trends against thermal time, during ripening of cultivar mixtures under well- and deficit-irrigation conditions.

| Moisture condition | Mixture treatments | $R^2$ | RMSE | Model P-value | Intercept | Slope | Pred.-Obs. thermal time to CCGR<sub>0</sub> (°Cd) | Pred. -Obs. days to CCGR<sub>0</sub> (day) |
| --- | --- | --- | --- | --- | --- | --- | --- | --- |
| Well irrigation | 1 | 0.868 | 0.034 | 0.000256 | -0.040 | 1.182 | 125.93 | 6.1 |
|  | 2 | 0.764 | 0.038 | 0.002068 | -0.017 | 0.937 | 122.16 | 5.9 |
|  | 3 | 0.915 | 0.023 | < 0.0001 | -0.078 | 1.466 | 134.14 | 6.5 |
|  | 4 | 0.869 | 0.028 | 0.000251 | -0.076 | 1.249 | 158.48 | 7.6 |
|  | 12 | 0.853 | 0.032 | 0.000372 | -0.045 | 1.044 | 138.56 | 6.7 |
|  | 13 | 0.879 | 0.031 | 0.000188 | -0.064 | 1.355 | 152.49 | 7.3 |
|  | 14 | 0.854 | 0.032 | 0.000372 | -0.038 | 1.131 | 137.61 | 6.6 |
|  | 23 | 0.811 | 0.032 | 0.000928 | -0.016 | 1.008 | 111.30 | 5.4 |
|  | 24 | 0.833 | 0.033 | 0.000589 | -0.044 | 1.260 | 146.78 | 7.1 |
|  | 34 | 0.899 | 0.027 | < 0.0001 | -0.080 | 1.476 | 145.01 | 7.0 |
|  | 123 | 0.822 | 0.035 | 0.000744 | -0.024 | 1.070 | 124.43 | 6.0 |
|  | 124 | 0.820 | 0.033 | 0.000784 | -0.022 | 1.038 | 145.72 | 7.0 |
|  | 134 | 0.893 | 0.027 | 0.000121 | -0.039 | 1.137 | 124.38 | 6.0 |
|  | 234 | 0.887 | 0.027 | 0.000149 | -0.058 | 1.216 | 141.82 | 6.8 |
|  | 1234 | 0.806 | 0.037 | 0.001024 | -0.013 | 1.055 | 128.15 | 6.2 |
|  | Mean | 0.852 | 0.031 | - | -0.044 | 1.175 | 135.80 | 6.537 |
| Deficit irrigation | 1 | 0.817 | 0.039 | 0.000832 | -0.013 | 1.011 | 134.16 | 6.5 |
|  | 2 | 0.746 | 0.040 | 0.002693 | -0.014 | 0.882 | 132.33 | 6.4 |
|  | 3 | 0.880 | 0.032 | 0.000184 | -0.077 | 1.510 | 147.58 | 7.1 |
|  | 4 | 0.842 | 0.034 | 0.000488 | -0.046 | 1.099 | 146.71 | 7.1 |
|  | 12 | 0.805 | 0.037 | 0.001035 | -0.017 | 0.893 | 137.00 | 6.6 |
|  | 13 | 0.900 | 0.029 | < 0.0001 | -0.093 | 1.467 | 159.31 | 7.7 |
|  | 14 | 0.828 | 0.039 | 0.000661 | -0.047 | 1.170 | 154.06 | 7.4 |
|  | 23 | 0.837 | 0.035 | 0.000550 | -0.057 | 1.145 | 145.88 | 7.0 |
|  | 24 | 0.854 | 0.033 | 0.000364 | -0.053 | 1.240 | 136.18 | 6.6 |
|  | 34 | 0.894 | 0.031 | 0.000116 | -0.084 | 1.446 | 145.44 | 7.0 |
|  | 123 | 0.862 | 0.032 | 0.000297 | -0.066 | 1.196 | 165.98 | 8.0 |
|  | 124 | 0.806 | 0.039 | 0.001024 | -0.027 | 1.057 | 133.51 | 6.4 |
|  | 134 | 0.820 | 0.042 | 0.000770 | -0.031 | 1.152 | 137.05 | 6.6 |
|  | 234 | 0.863 | 0.033 | 0.000292 | -0.060 | 1.219 | 139.21 | 6.7 |
|  | 1234 | 0.854 | 0.037 | 0.000364 | -0.058 | 1.220 | 136.89 | 6.6 |
|  | Mean | 0.841 | 0.035 | - | -0.050 | 1.180 | 143.42 | 6.9 |

Mixture treatments include the monocultures and mixtures of the four early- to middle-ripening wheat cultivars. Accordingly, each digit stands for a single cultivar included in the mixture, so the treatment 1234 shows the 4-component mixtures of cultivars 1,2,3 and 4. $R^2$ , $RMSE$ , $Model\ P\text{-}value$ , $Intercept$ , and $Slope$ are the parameters of the linear regression between the predicted and observed data. CCGR<sub>0</sub> is the time (and/or thermal time) when CCGR becomes zero (at ripening). " $Pred. -Obs.$ " shows the difference between the predicted and observed values. Number of days in the last column are estimated by dividing the corresponding thermal times (in the adjacent column) by the averaged diurnal thermal times of the final 10 days.

**Figure 4.**
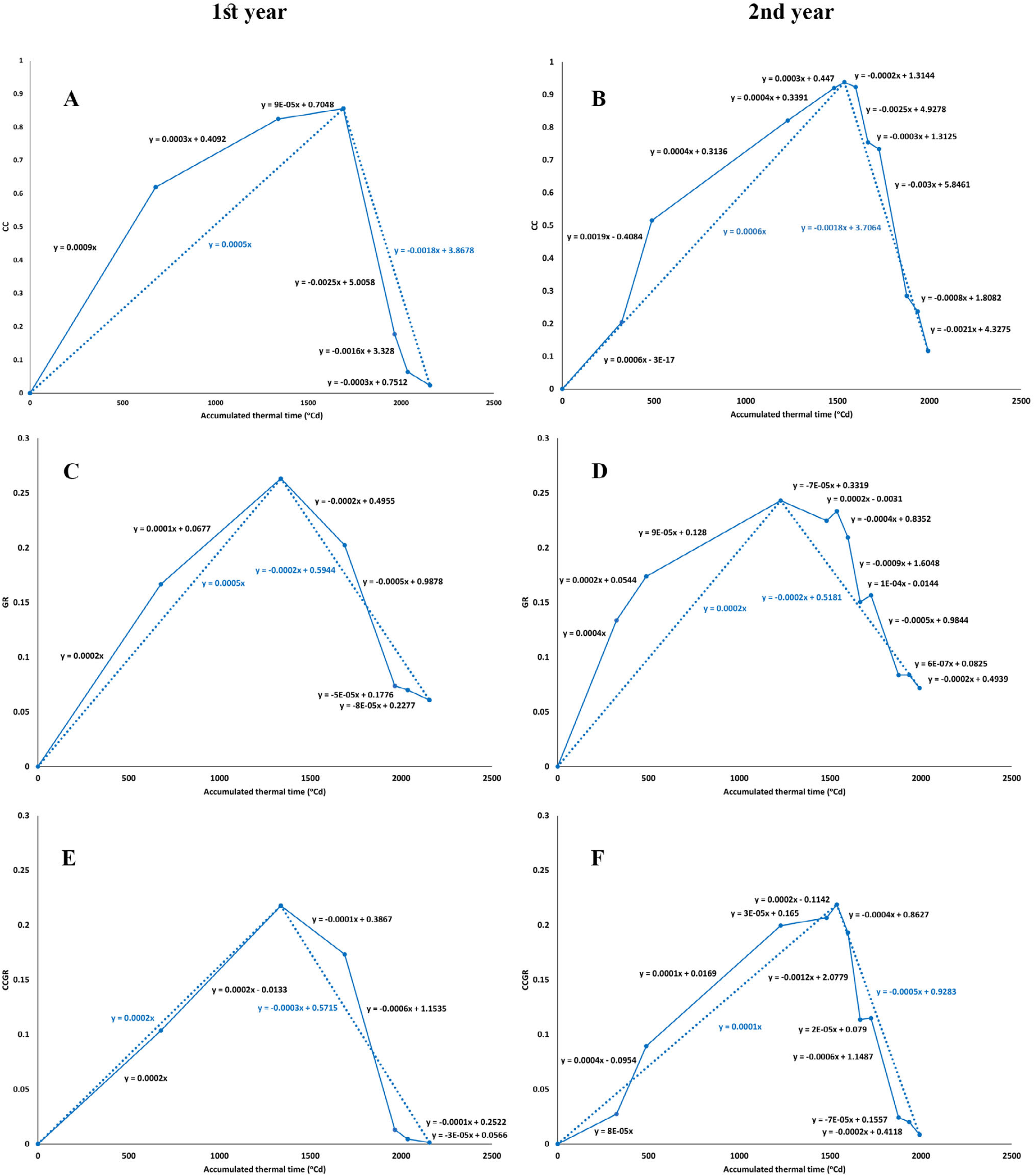
The split linear trends of the image-derived criteria of CC, GR, and CCGR throughout the 1 st season, calculated for the monoculture of cultivar 1 under well-irrigation conditions. The dotted lines with positive and negative slopes indicate the overall linear trends from sowing to the observed peaks, and then from the maximum values to the least recorded ones at the late season, respectively (the equations with light colors represent the dotted lines).

As shown in Fig. S9, the CC linear declining trends of the mixture treatments (during ripening, from CC_max_ to CC_0_) were divergent towards the end of season, under well-irrigation conditions; while the same trends were more parallel under the deficit-irrigation condition. It is notable that this linear trend (CC_max_ to CC_0_) could not predict the ripening time (CC_0_), and was not comparable with the accuracy of the binomial equations described before. Tables S7 to S9 represent the effect of mixture and irrigation treatments on the linear trends of the image-derived indices, and the results of mean comparisons over the two years. Obviously, the significant effects of the mixture or irrigation treatments, and also their interaction on the linear trends were frequent (Table S7), which may indicate the potential and sensibility of the split linear trends in detecting the different temporal trends among treatments.

Another advantage of using the split linear trends in evaluating the short-term variations and/or fluctuations of the image-derived indices, relying on the variation between the consecutive points (imaging dates) is shown in Fig. S10. In this figure, the results of monocultures in the second year of the study were represented (since the imaging events were more frequent in this year, so a higher temporal resolution for detecting the minor variations was provided). The linear slope of CC between the second and third imaging dates had increased steeply, compared with the previous and the preceding trends, and despite the reduced thermal times (in the winter). This trend coincided with the early-to middle-tillering growth stage and the respective canopy coverage development. Oppositely, in the same period, the previously sharp linear slopes of GR declined gradually, as an initiation for the meantime trend of the declining slopes till reaching the maximum GR.

Furthermore, two irrigation events during the ripening, made two sets of minor fluctuations in overall declining trends of the either image-derived indices, irrespective of the irrigation treatment. Interestingly, as the CC fluctuations indicate, the responses of the cultivars to the irrigation in the late season, were in the order of their ripening rate; so the 1^st^ and 2^nd^ cultivars were less influenced, while the CC of the 3^rd^ and 4^th^ cultivars were even increased, in contrast to the general direction of ripening. Such increasing is expected to be the consequence of altering leaf angels after irrigation, in the more stay-green canopies. Similar responses are also evident for the GR and CCGR indices. In general, these results show that each irrigation event had temporarily postponed the overall ripening trend for several days; the process which was associated with both green surface quantity and quality.

### Predicting the image-derived indices of the mixtures, based on the corresponding monocultures

A reasonably estimation approach suggested for evaluating cultivar mixtures, is predicting the intended factors using the averaged amounts of the monocultures included in the mixtures (Finckh and Mundt, 1992; Mille *et al*., 2006). Figure 5 indicates the diversion of the observed diurnal CC, GR, and CCGR from the predicted values estimated based on the averaged amounts of the respective monocultures, under various irrigation conditions during the two seasons. Despite the differences between the results of the 1^st^ and 2^nd^ year, as the variation ranges were narrower in the latter, the ranges were increasingly enhanced towards the end of season, i.e. during ripening (Figure 5). Among the indices, GR and CCGR had the lowest and highest ranges of variations, respectively, regardless of the year and irrigation treatment. Generally, the amounts of variation seem to be considerably high, as for instance, the biases up to almost ±60% and ±40% were frequent in the 1^st^ and 2^nd^ years, respectively. An agronomic implication of such comparisons are represents in the Figure S11, where the two-component mixture of the 1^st^ and 4^th^ cultivars under deficit-irrigation condition (treatment 14) were compared with the corresponding monocultures. The mixture had a higher degree of stay-green than either of the monocultures, (Fig. S11; see the images and diagrams), as CC and CCGR values of this mixture were approximately 3 times higher than the values in the more late-ripening monoculture, while the differences between the GR amounts were lower. Although particularly under the terminal water stress conditions, the resulted higher stay-green characteristic may be a valuable advantageous, the related mechanisms or reasons are outside the scope of this paper. It also should be noted that such differences among the canopies may be not necessarily such apparent in the images, so that the comparisons would be possible only by utilizing the quantified indices.

**Figure 5.**
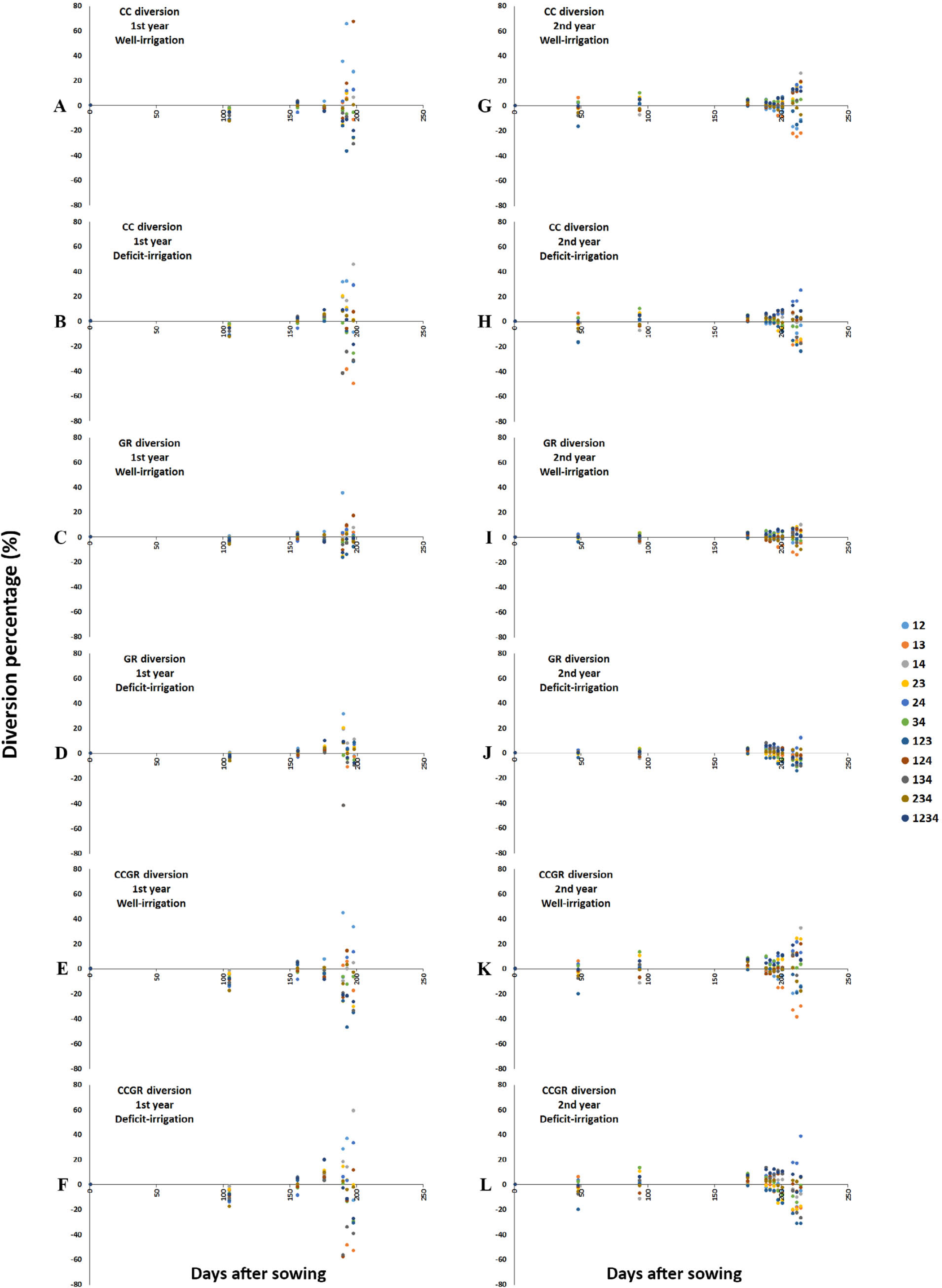
The diversion of the observed CC, GR, and CCGR values of the mixtures over the season, from the predicted values calculated as the averaged amounts of the corresponding monocultures (e.g. in the part A, the CC value of the mixture 124 is more than 60% higher than the average amount of the monocultures 1, 2, and 4 under the same conditions. Before the late season, the color circles are mostly overlapped around the zero line.

Using the similar approach for evaluating the mixtures’ diversions from the average of the corresponding monocultures, the ripening date (i.e. CC_0_ calculated based on the binomial model) was also considered, besides the diurnal values. As Fig. S12 indicates, despite for the diurnal values described before, the maximum diversions were less than 2 dates, irrespective the year or irrigation conditions, though, the biases were even relatively lower under the post-anthesis deficit irrigation.

### The relationship between CC and GR

The relationship between CC and GR (i.e. the quantitative parameters represents quantity and quality of the green canopy), was also evaluated (Fig. 6, 7, and Table S10). In both years, they did not show any significant relationship unless at a particular stage during ripening, when the correlation (and also regression) parameters raised to a significant peak. This peak was recorded at 192 DAS in the first year, and jointly at 200 (for both well- and deficit-irrigation) and 208 DAS (under well-irrigation) in the second year. Although based on the thermal time or days after sowing, the phenomenon occurred at different times (in both growing seasons), the conventional phenological stages were almost identical; which seems to be the best criteria for determining the time of reaching the highest correlation between CC and GR. The respective growing stage was at soft dough in the most early-ripening cultivar, synchronized with the milk stage of the most late-ripening one, in both years (Fig. S13, see the cultivars’ growing stages at 192 and 200 DAS in the 1^st^ and 2^nd^ year, respectively).

**Figure 6.**
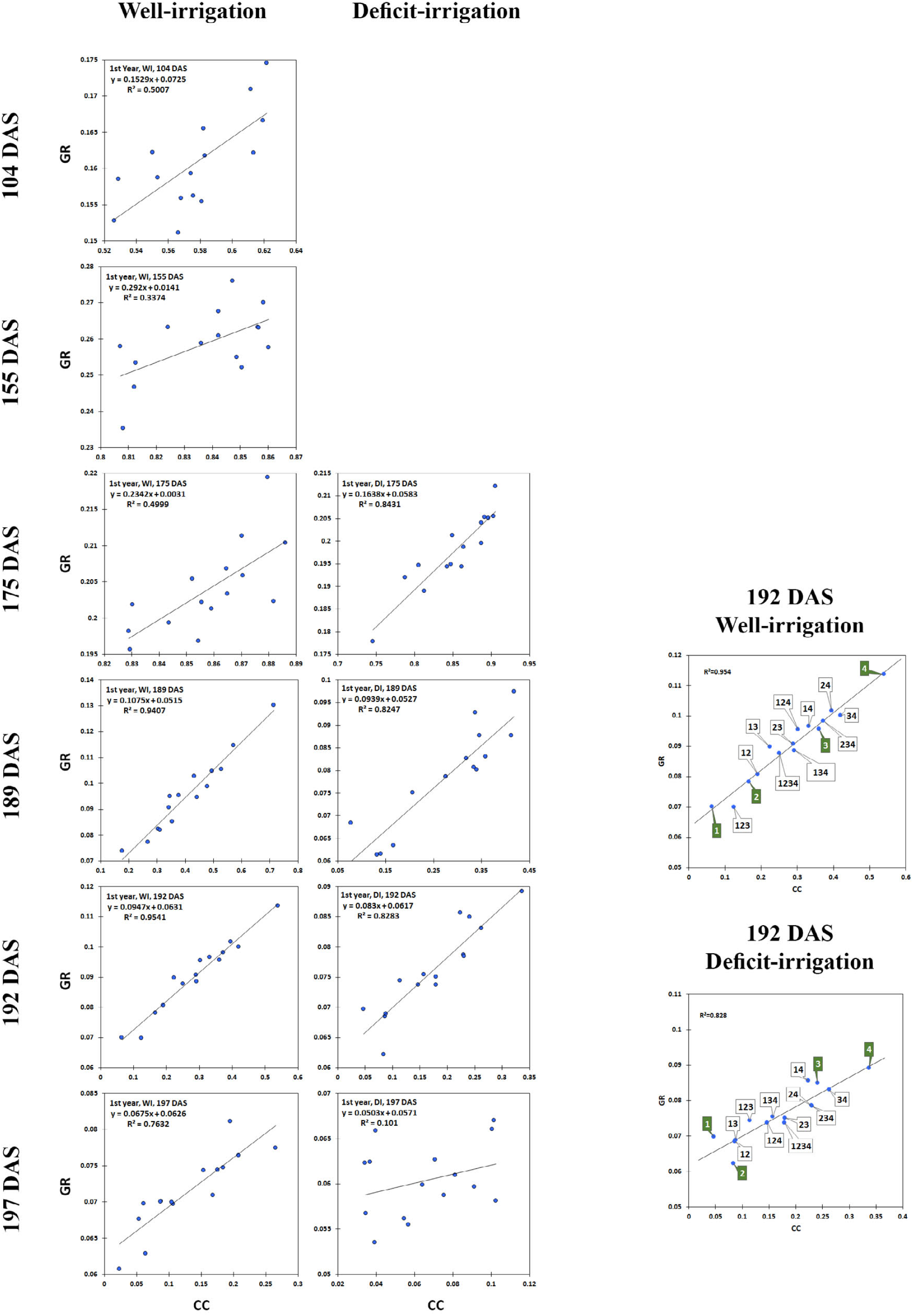
The relationship between CC and GR under well- and post-anthesis deficit-irrigation conditions during the lit year. DAS: days after sowing. The two detailed charts at the right side of the figure, represent the ranking of monocultures and mixtures, in the date with the highest relationship between CC and GR.

**Figure 7.**
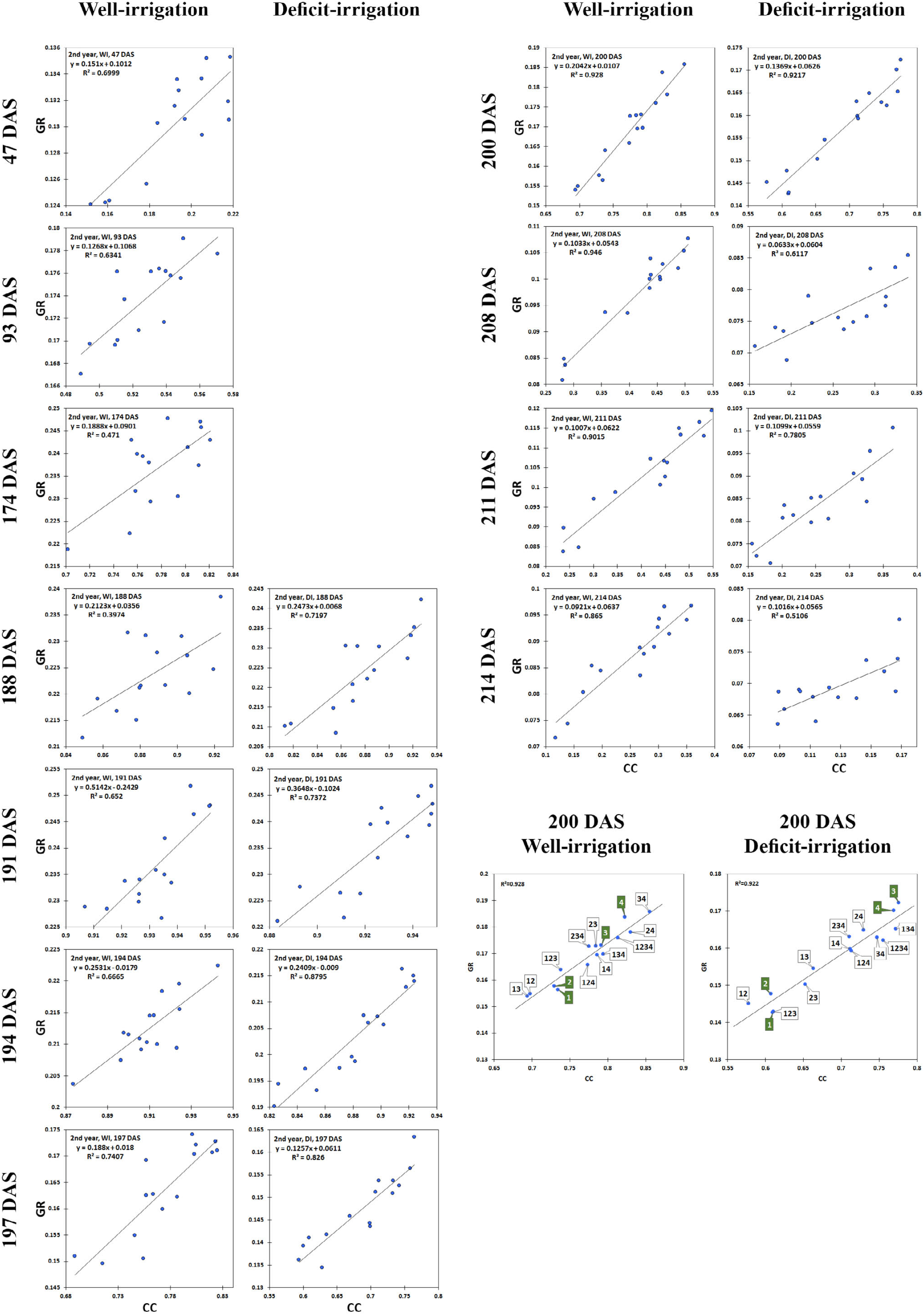
The relationship between CC and GR under well- and post-anthesis deficit-irrigation conditions during the 2nd year. DAS: days after sowing. The two detailed charts at the end of the right column, represent the ranking of monocultures and mixtures, in the date with the highest relationship between CC and GR.

It is obvious that at the time when CC and GR show the strongest relationship, the cultivars with the highest values of CC had also the highest amounts of GR, and vice versa. Since having the maximum CC and GR simultaneously is considered as a physiological advantage, particularly late at the season, it might provide a potential opportunity for comparing the mixtures (and also cultivars) arranged in a reasonable linear low-to-high order. Therefore, seeking more evidences for understanding the related implications and consequences, the relative positions of the treatments - especially the monocultures-in the mentioned line were compared with some other measured physiological trends including grain yield and yield components (data not shown). However, no considerable similarities were found, except for the order of ripening, which may explain the condition clearly (the pairs of detailed regression charts in Fig. 6 and 7). Obviously, the arrangement of mixture treatments (including monocultures and mixtures) with the order of least-to-highest CC and GR values followed the ripening patterns of the 1^st^, 2^nd^, 3^rd^, and 4^th^ cultivars, respectively. For instance, the 4^th^ cultivar had the highest rank as the most late-ripening one, while the early ripening cultivars along with their respective mixtures were in the lowest rank. More research is needed to understand whether the reported observations about the relationship between CC and GR, are limited to the situations of the present study or may be extrapolated to other conditions and genotypes. Besides, for more information about the comparative trends of the image-derived indices and conventional phenology, (see Figure S13) which represents the brief timelines of the events during the ripening of the 4 monocultures evaluated.

## Discussion

The ripening trends of the mixtures were monitored and evaluated quantitatively using the declining trends of three image-derived indices including: (i) CC, which is a well-known index associated to the quantity of the green canopy cover; (ii) GR, a modified index related to the quality of the reflected light from green surfaces; and (iii) the new introduced index of CCGR, which can show the quantity and quality of the green surfaces of the canopy, integratively. Besides having simple formulae, as indicated by the statistical analyses (e.g. on the diurnal values), the indices seemed to show acceptable degrees of sensibility to the crop variations, high consistency with the visual observations (or images), and were straightforward and easily interpretable; the beneficial properties which may maintain them among the efficacious choices for tracking the crop temporal trends, depended on the objectives.

During ripening, the binomial equations showed the best fits on the declining trends of the image-derived indices regressed on the accumulated thermal times, though their rates were different. Accordingly, CC values became zero late in the season earlier than the other two indices, therefore CC_0_ was selected as the criterion for ripening time in the diverse canopies of mixture treatments. The binomial trends of the indices could distinguish between the ripening of the 4 cultivars, and revealed that the post-anthesis deficit-irrigation had shorten both the ripening period and the diversity among the cultivars’ ripening time, regardless of the year. such findings might not have been detected in the heterogeneous mixtures, unless using the quantitative criteria. The results of model validations also showed the predictability of the indices based on the accumulated thermal times, -and also with a caution-on the GDDs. Correspondingly, the biases between the predicted and observed terminal zero values of the indices (the ATT/ or GDD at which the index quantity had been extinct completely) as the image-based events were as large as several days (in average, 5.2 to 6.2 for CC, 9.4 to 10.2 for GR, and 6.5–6.9 for CCGR, under well-and deficit-irrigation, respectively). Such amounts seem to be acceptable, respective to some other reports about using the remote sensing indices to predict a quantified event (e.g. Johnen *et al*., 2012). Moreover, when the purpose of the analyses is calculating the actual rates of the increasing or decreasing trends rather than their model-based estimated values, the observed points should also be utilized; as was carried out for calculating the average rates of the CC values from sowing to the maximum recorded peak, or from the peak to the terminal low extremes. However, for ensuring the capturing of the real peaks, or achieving them with the least diversions, the imaging events is required to be frequent enough. An example of such requirement may be the case for calculating the positive or negative linear slopes using the observed CC_max_ in the first year, which seems would lead to results with higher consistencies between the two years, if there were more observations around the current recorded imaging date (i.e. CC_max_ in the first year).

In addition to utilizing the overall directions of the variations for evaluating the mid- to long-term trends of the canopy, the split linear trends were also assessed in order to interpret the temporary (short-term) fluctuations of the indices values (similar to Magney *et al*., 2016). Clearly, in the first strategy i.e. focusing on the long-term (binomial or linear) trends, the entire trend fitted to the data points is considered as the first priority, and the individual points (observations) and/or the temporary fluctuations might be ignored; while in the second strategy (split linear trends), even the pairs of points and minor fluctuation are taken into considerations. Among the instances, are determination of the ripening time based on the entire trend line, versus distinguishing the effect of irrigation events by concentrating on the minor index fluctuations.

Utilizing the image-derived indices, some other agronomic assessments were also carried out, including predicting the mixtures behavior based on the included monocultures, and evaluating the relationship between CC and GR, as the quantitative indices for quantity and quality of the canopy green coverage. The results showed that at least in the conditions of the present study, the diurnal quantities of the evaluated indices in the mixtures tended to be diverted increasingly from the averaged values of the respective monocultures, towards the late season. The high diversions at the late season may provide considerable evidences for either synergetic or antagonistic inter-cultivar relationships within the mixtures, which in the first case, can make potential opportunities for selecting the beneficial cultivar mixtures e.g. in order to improving the canopy stay-green, particularly under the stressful conditions late in the season. As described before, the situation may be influenced by the year and water stress. Despite the frequent diversions between the diurnal values of the mixtures and monocultures, the predicted mixtures’ ripening times (CC_max_) based on the binomial trends of the corresponding monocultures, showed lower errors (less than 1 or 2 days) compared with the observed ripening dates.

The linear relationship between CC and GR was weak, except in a critical growth stage in both seasons, when they showed high correlations. In the respected imaging dates, the relative ranking of the mixture treatments (especially monocultures) in having the highest amounts of CC and GR followed the ripening patterns of the cultivars included.

As evidenced in the present study, common digital images may represent an extremely informative source for studying the canopy temporal trends quantitatively, in the light of reasonable indices and computation methods. However, no successful approach would be created unless the preconditions are met. Primarily, these quantitative remote sensing-based indices should provide an appropriate reflection of the crop respond to the progress of the parameter taken as the driving factor of the phenological dynamics (i.e. time, thermal time, cumulative growth degree days, etc.). In other words, the candidate indices should be sensible enough to the crop growth and development. Accordingly, as another preference, the candidate indices are also expected to be predictable enough to be used as alternatives for the conventional phenological events where needed (e.g. in the nowadays crop models). Furthermore, the most consistent data processing method with the objectives should be selected and utilized. For example, non-phenological fluctuations of the amounts should be distinguished and -depended on the purposes-either be included in or excluded from the calculations. The non-phenological variations are expected to be temporary (e.g. in the scale of hours to several days) and may be associated with field management including irrigation or fertilization, biotic or abiotic stresses, and the data acquisition practices (e.g. the time of imaging or sensor readings). Therefore, additional information i.e. about the occurred fluctuations during the cropping season may be required in practice, for interpretation of the variations in the index’s trend; even including the conventional phenology, which was excluded from the main computational stream. Indeed, the conventional phenology may be useful for casual validation and checking that whether a given data point may be included in the evaluations or not. Further, it may provide the key for understanding and interpreting the apparently incompatible coincident trends, e.g. as mentioned before, the increasing trend of CC despite the declined diurnal degree days was explained by the crop growth stage (tillering) based on the conventional phenology and related biological facts (low temperatures requirement for tillering). Therefore, if utilized and integrated appropriately, it is expected that the novel independent quantitative frameworks and the conventional phenology may be synergetic for describing the temporal crop trends, efficiently.

## Conclusion

In the present study, the option of monitoring and quantifying the ripening trends in the heterogeneous stands of wheat cultivar mixtures was evaluated using a commercial digital camera, independent of the conventional phenology. For this purpose, three simple image-derived indices, including the well-known canopy cover (CC), and the modified of G-R (i.e. GR) indices were employed as quantitative criteria for quantity and quality of the green surfaces in the canopy, and also the novel index of CCGR was introduced for analyzing the quality of reflected light (GR) from the green surfaces per unit area. The results showed that the different quantities of the indices regressed against thermal time may be taken as the new phenological events, depended on the purposes. Accordingly, the binomial trends showed the best fit to the declining trend of either index during ripening; by which, it was also shown that the utilized image-derived indices may be predictable based on the accumulated thermal time. Besides, some agronomic aspects were described using the various estimation methods based on the indices, including: (i) the post-anthesis deficit irrigation accelerated the ripening, and reduced the diversity of ripening dates among the cultivar mixtures; (ii) the short-term fluctuations in the values of the image indices revealed by the split linear trends, could reflect the irrigation events and their different comparative effects on the early- to middle-ripening cultivars; (iii) the relationship between CC and GR was not strong unless at the soft-dough and milk stage of the early- to the most late-ripening cultivars, respectively. The suggested indices appeared to have the potential use in developing independent quantitative frameworks, parallel to the conventional qualitative phenology, though they may contribute integratively to the interpretation of the temporal crop trends.

## Acknowledgement

The authors wish to thank Shiraz University for providing field experiment facilities.

---

**Supplementary Figure S1.**
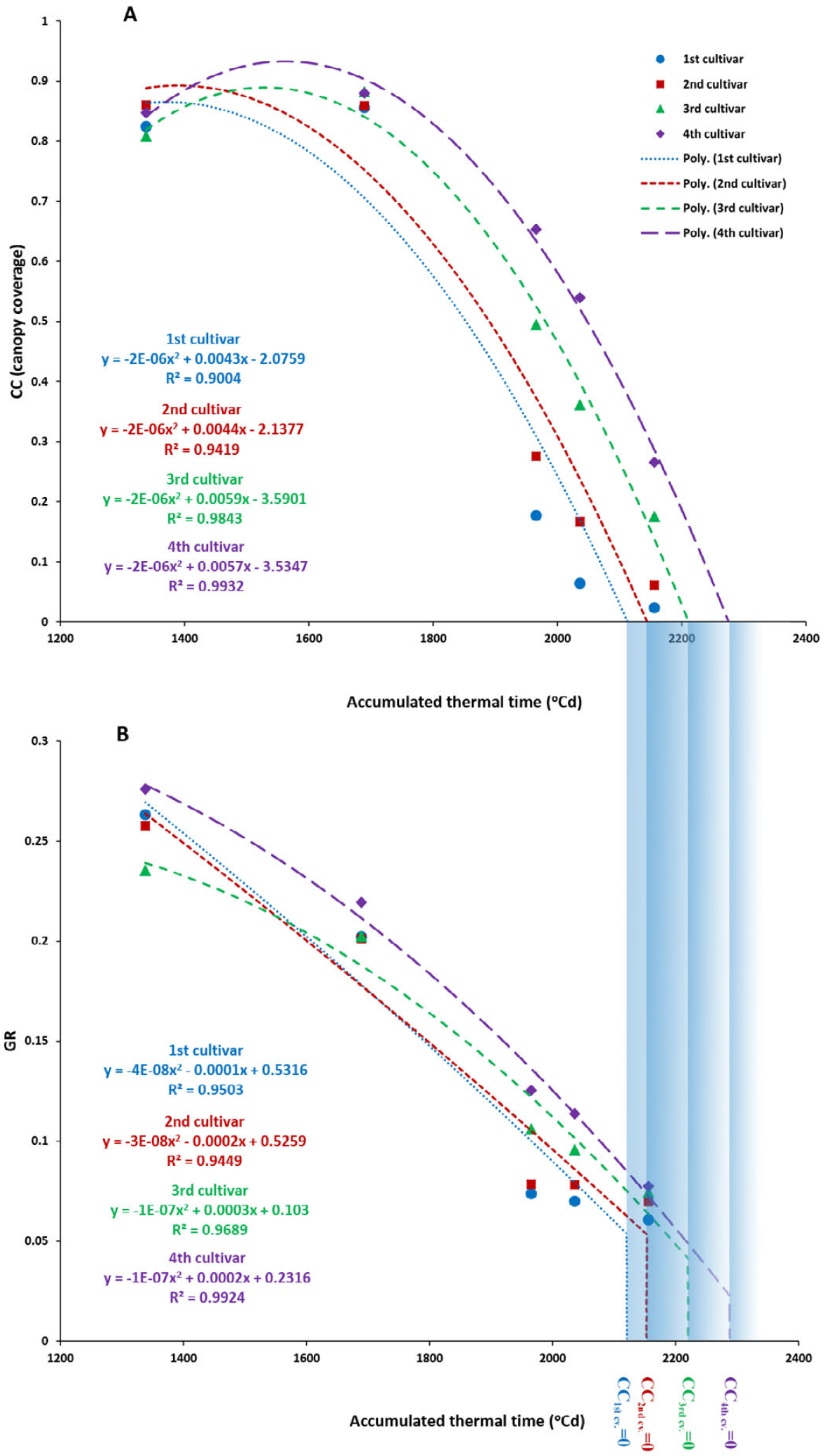
The predicted CC value of either cultivar reach to zero earlier than the predicted GR based on the binomial models.

**Supplementary Figure S2.**
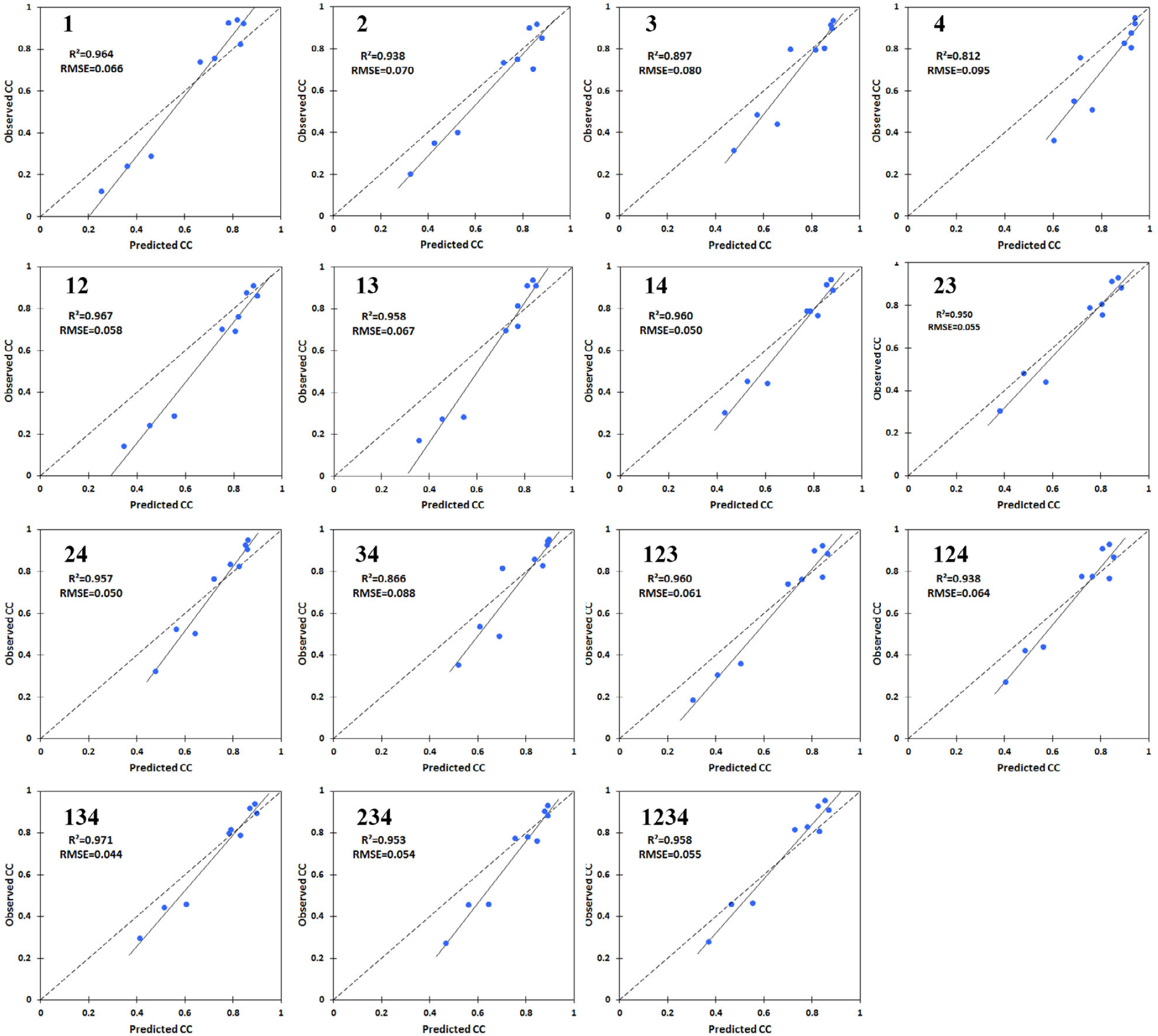
The relationship between the predicted and observed CC (canopy cover) values of the mixture treatments under well-irrigation conditions. Each digit in the treatment titles show a cultivar included in the mixture. 1, 2, 3, and 4 are the four early- to middle ripening wheat cultivars, respectively.

**Supplementary Figure S3.**
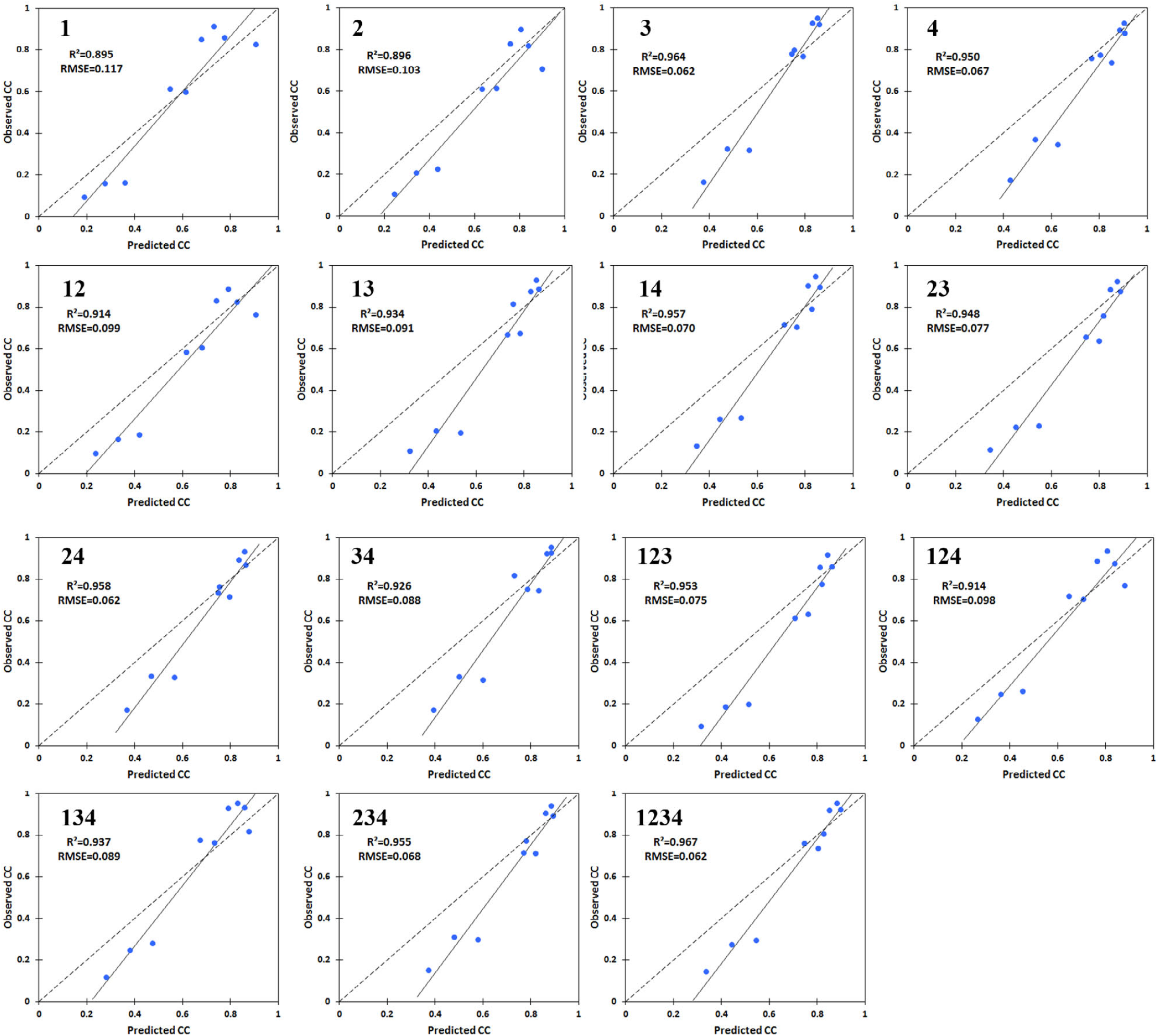
The relationship between the predicted and observed CC (canopy cover) values of the mixture treatments under post-anthesis dificit-irrigation conditions. Each digit in the treatment titles show a cultivar included in the mixture. 1, 2, 3, and 4 are the four early- to middle ripening wheat cultivars, respectively.

**Supplementary Figure S4.**
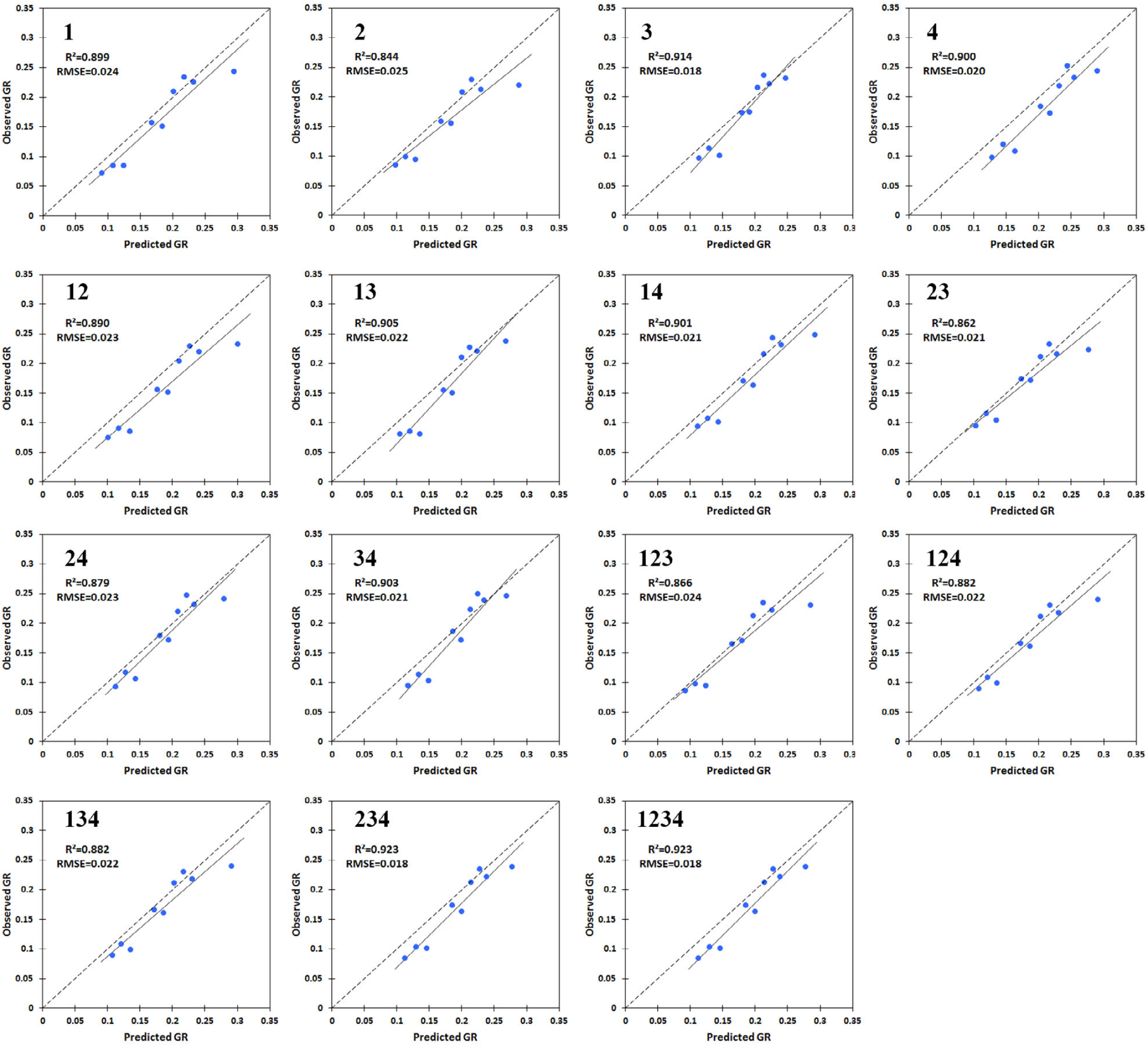
The relationship between the predicted and observed GR values of the mixture treatments under well-irrigation conditions. Each digit in the treatment titles show a cultivar included in the mixture. 1, 2, 3, and 4 are the four early- to middle ripening wheat cultivars, respectively.

**Supplementary Figure S5.**
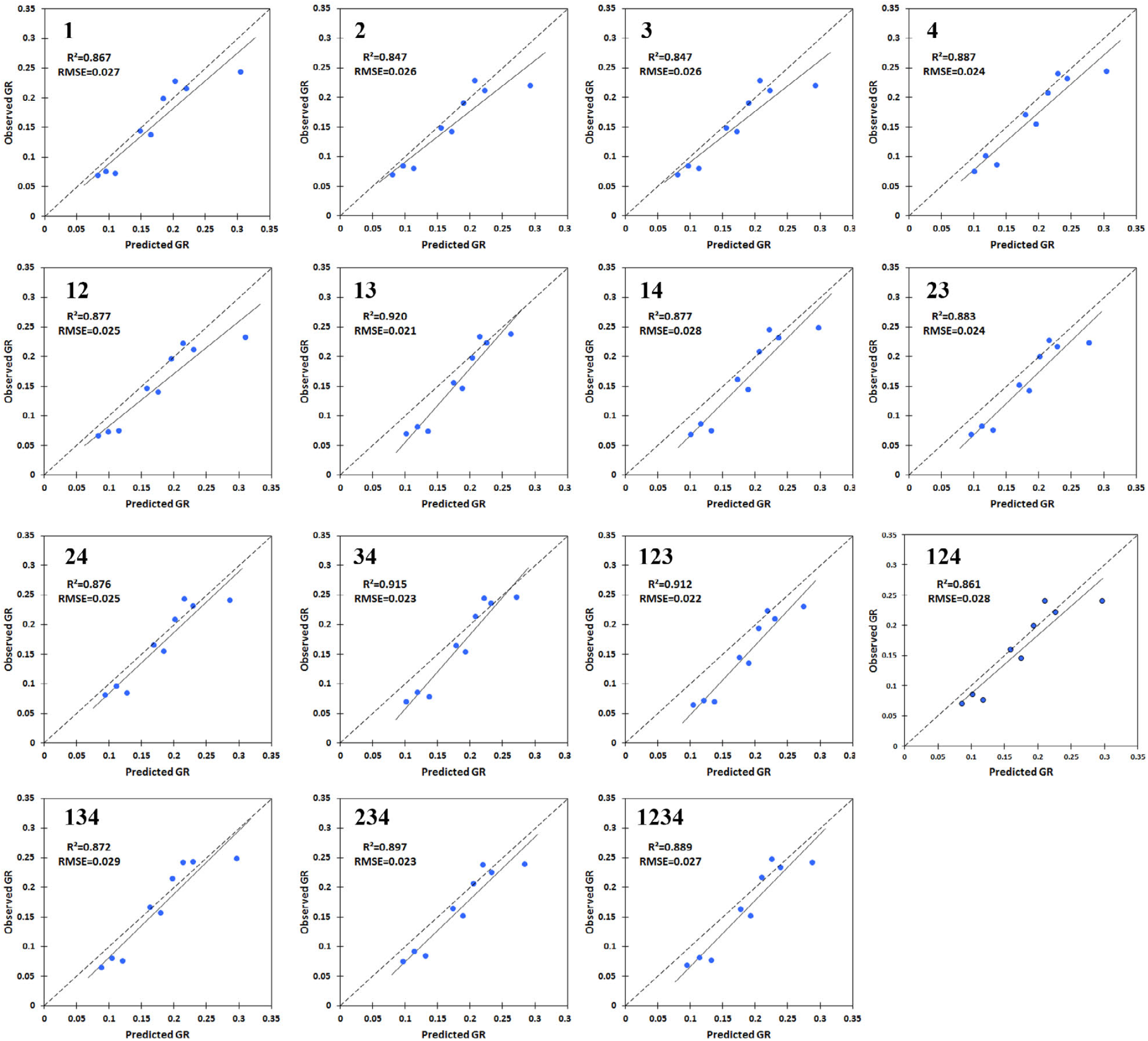
The relationship between the predicted and observed GR values of the mixture treatments under post-anthesis deficit-irrigation conditions. Each digit in the treatment titles show a cultivar included in the mixture. 1, 2, 3, and 4 are the four early- to middle ripening wheat cultivars, respectively.

**Supplementary Figure S6.**
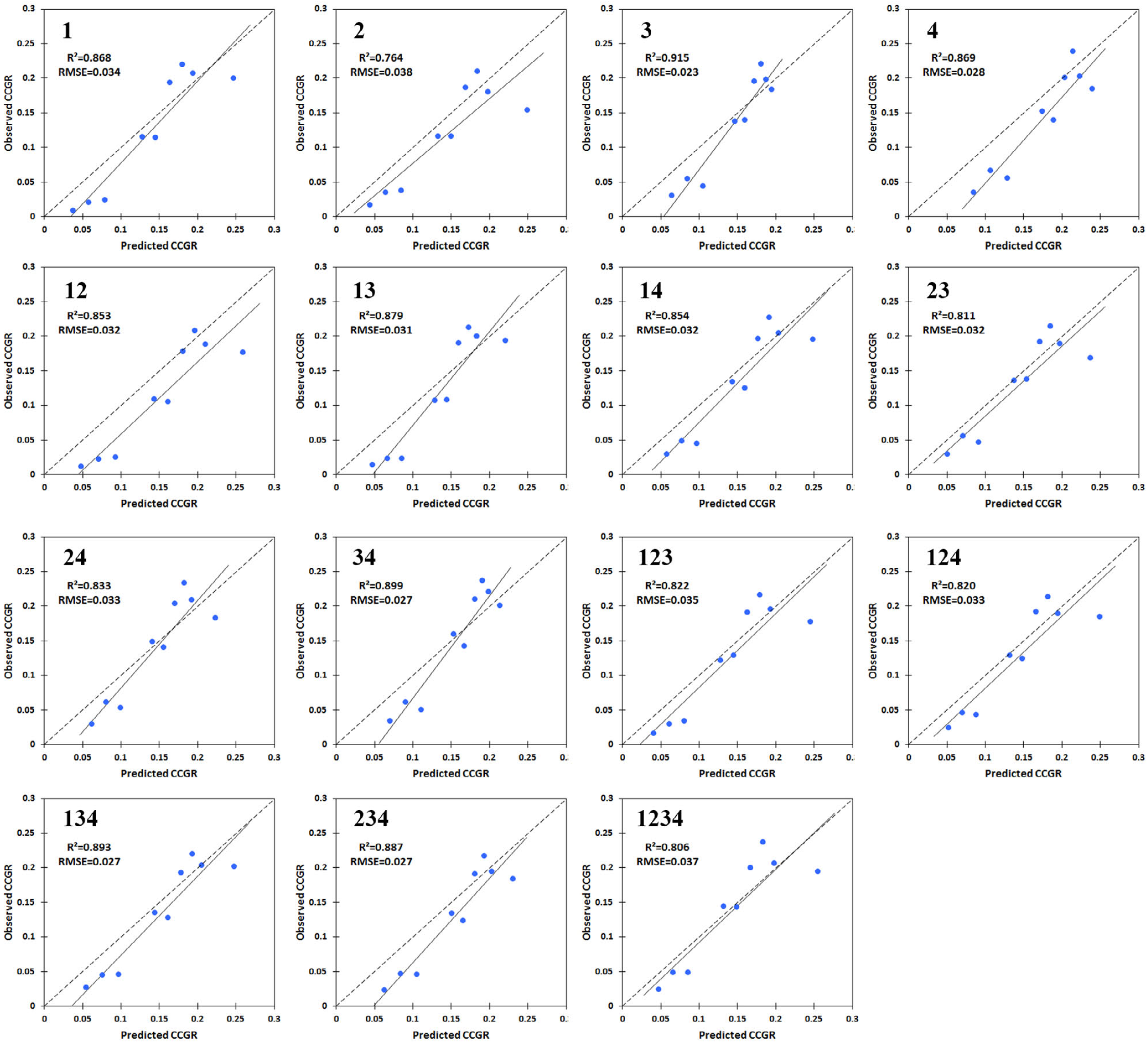
The relationship between the predicted and observed CCGR values of the mixture treatments under well-irrigation conditions. Each digit in the treatment titles show a cultivar included in the mixture. 1, 2, 3, and 4 are the four early- to middle ripening wheat cultivars, respectively.

**Supplementary Figure S7.**
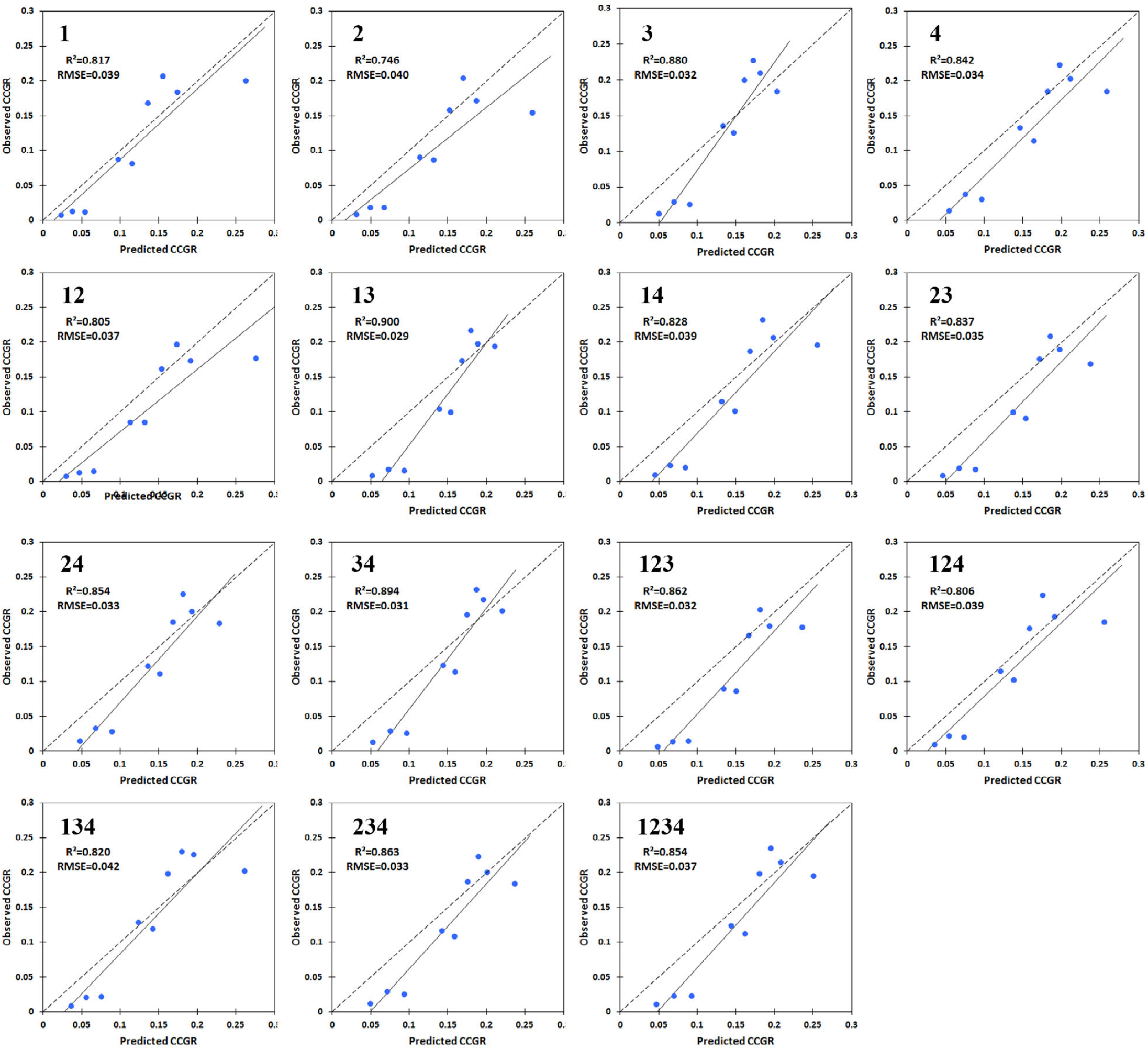
The relationship between the predicted and observed CCGR values of the mixture treatments under post-anthesis deficit-irrigation conditions. Each digit in the treatment titles show a cultivar included in the mixture. 1, 2, 3, and 4 are the four early- to middle ripening wheat cultivars, respectively.

**Supplementary Figure S8.**
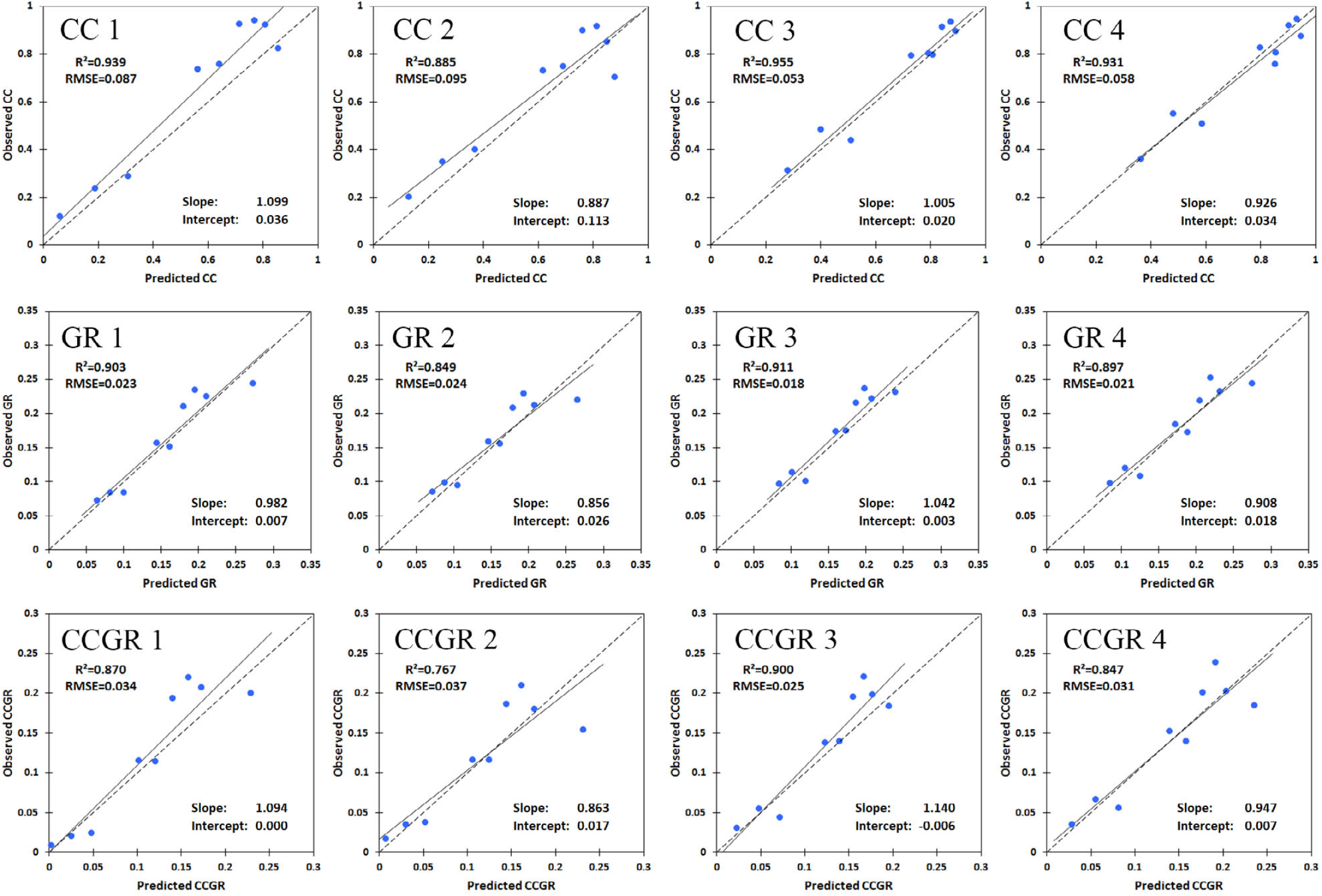
The relationship between the predicted and observed values of the image-derived criteria (calculated based on GDD-growing degree days-) for monocultures, under well-irrigation conditions. 1, 2, 3, and 4 are the four early- to middle ripening wheat cultivars, respectively.

**Supplementary Figure S9.**
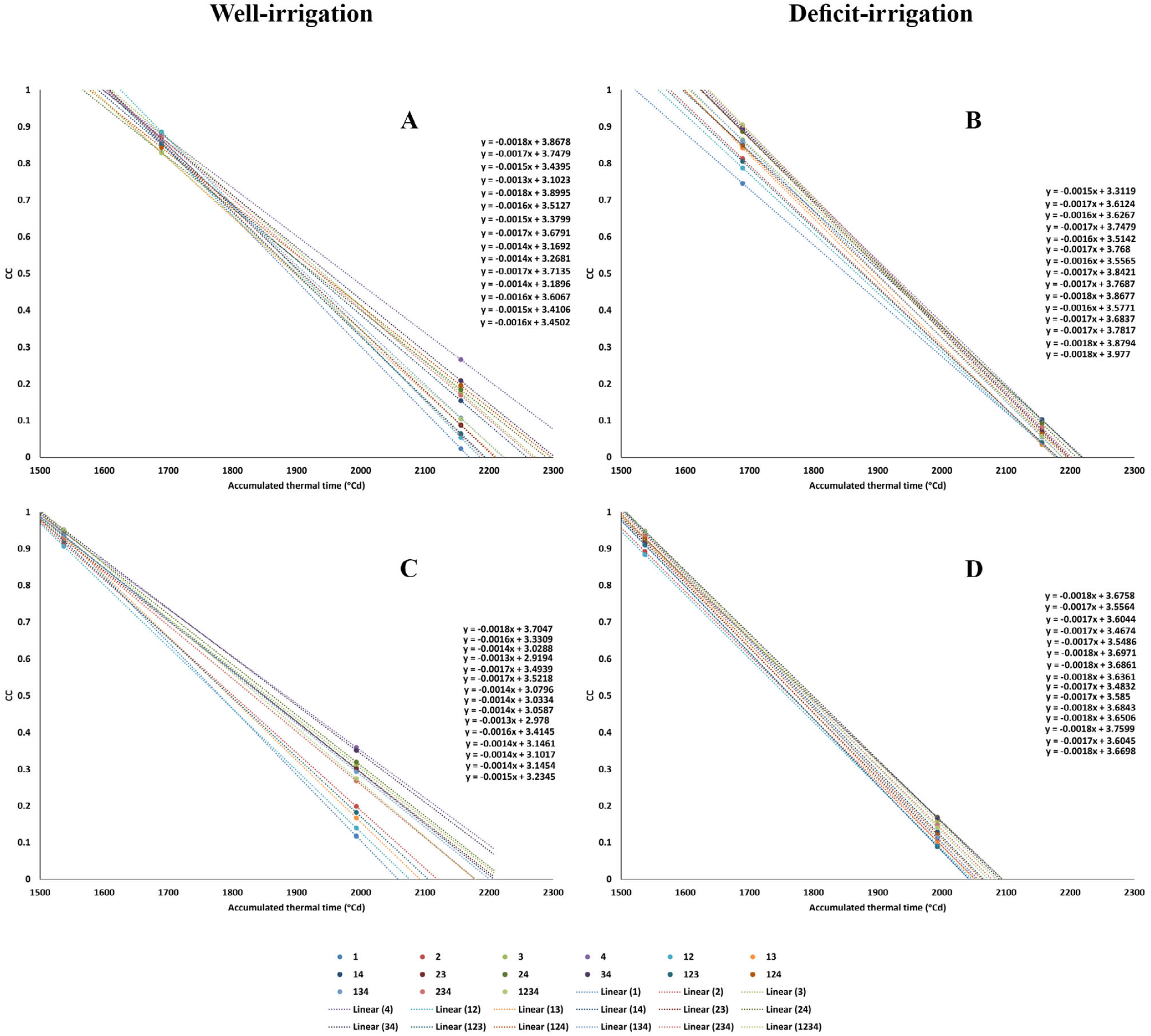
The linear declining trends of CC from the observed seasonal peak (maximum amount) to the last imaging date in the monocultures and mixtures of the 4 early- to middle-ripening winter wheat cultivars; **(A)** and **(B)**: in the last year, **(C)** and **(D)**: in the 2nd year. The equations arranged from top to down, represent the trends of the treatments in the order below: 1, 2, 3, 4, 12, 13, 14, 23, 24, 34, 123, 124, 134, 234, 1234; where 1, 2, 3, and 4 are the monocultures of the early to middle ripening cultivars, respectively, and the other treatments are the mixtures included these cultivars, e.g. the treatment 1234 is the 4-component mixture of the 4 cultivars. Largely due to the relatively higher diversity in the CC values of the mixture treatments at the lait imaging date under well-irrigation conditions of both season, the linear trends are divergent towards the late season; while, under the post-anthesis deficit-irrigation, the linear trends are almost parallel.

**Supplementary S10.**
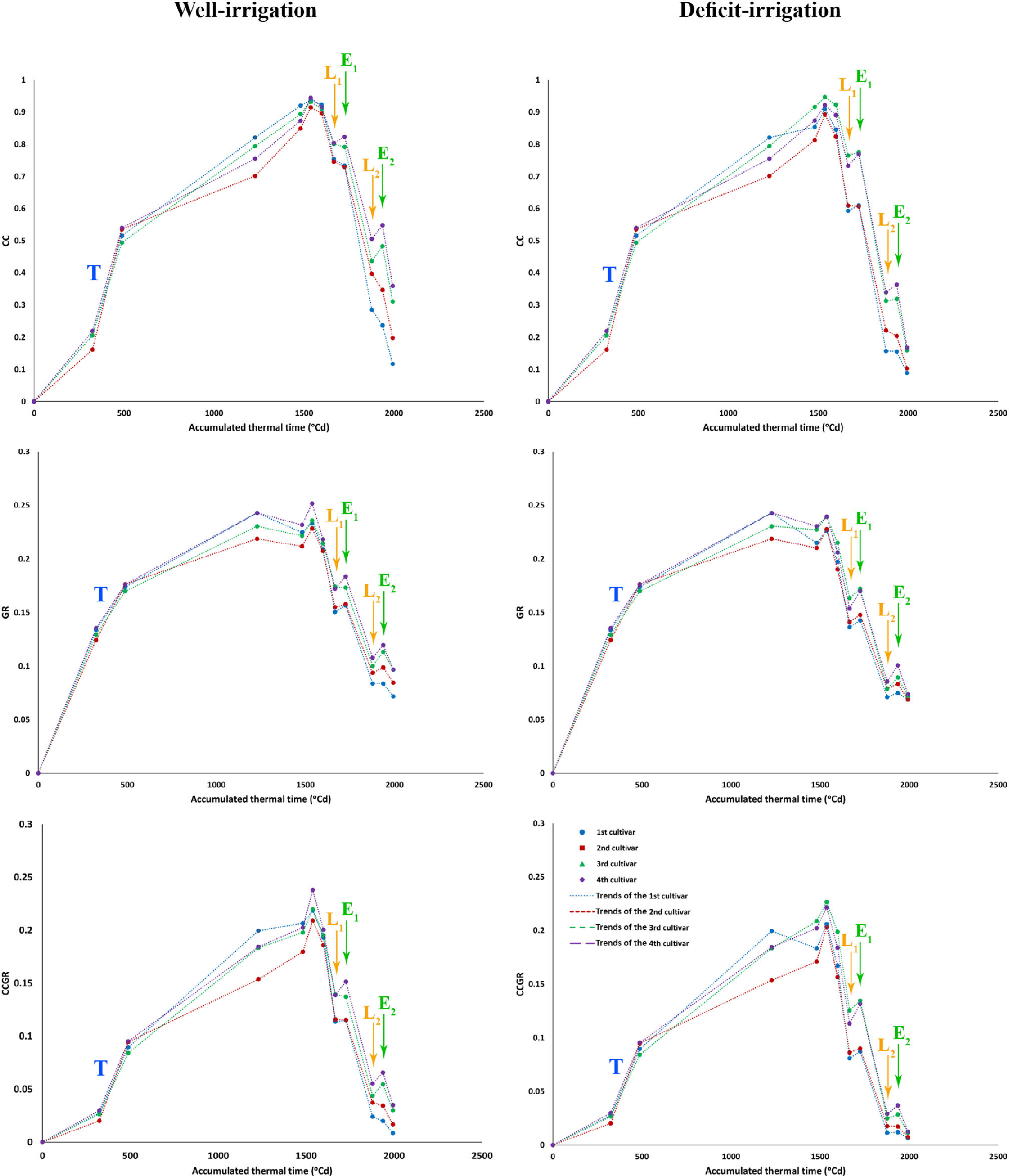
The short-term variations of the image-derived criteria under well- and deficit-irrigation conditions in the monocultures, during the 2^nd^ season. The l^st^ 2^nd^, 3^rd^, and 4^th^ cultivars are the early- to middle-ripening cultivars, respectively. “T” shows the trends during the early tillering growth stage, which have relatively steep increasing slopes in the cases of CC and CCGR (obviously despite the low diurnal temperatures in this period); while the “T” trends for GR have lower slopes. “L” and “E” indicate the imaging events, before and after each irrigation during the ripening, i.e. late and early in the irrigation intervals, respectively. The corresponding imaging dates were as follows: **L_1_**: 197 DAS (1 DBI), **E_1_**: 200 DAS (2 DAI), **L_2_**: 208 DAS (0 DBI, i.e. exactly before irrigation), and **E_2_**: 211 DAS (3 DAI); where DAS, DBI, and DAI are: days after sowing, days before irrigation, and days after irrigation, respectively. The irrigation intervals were 10 days.

**Supplementary Figure S11.**
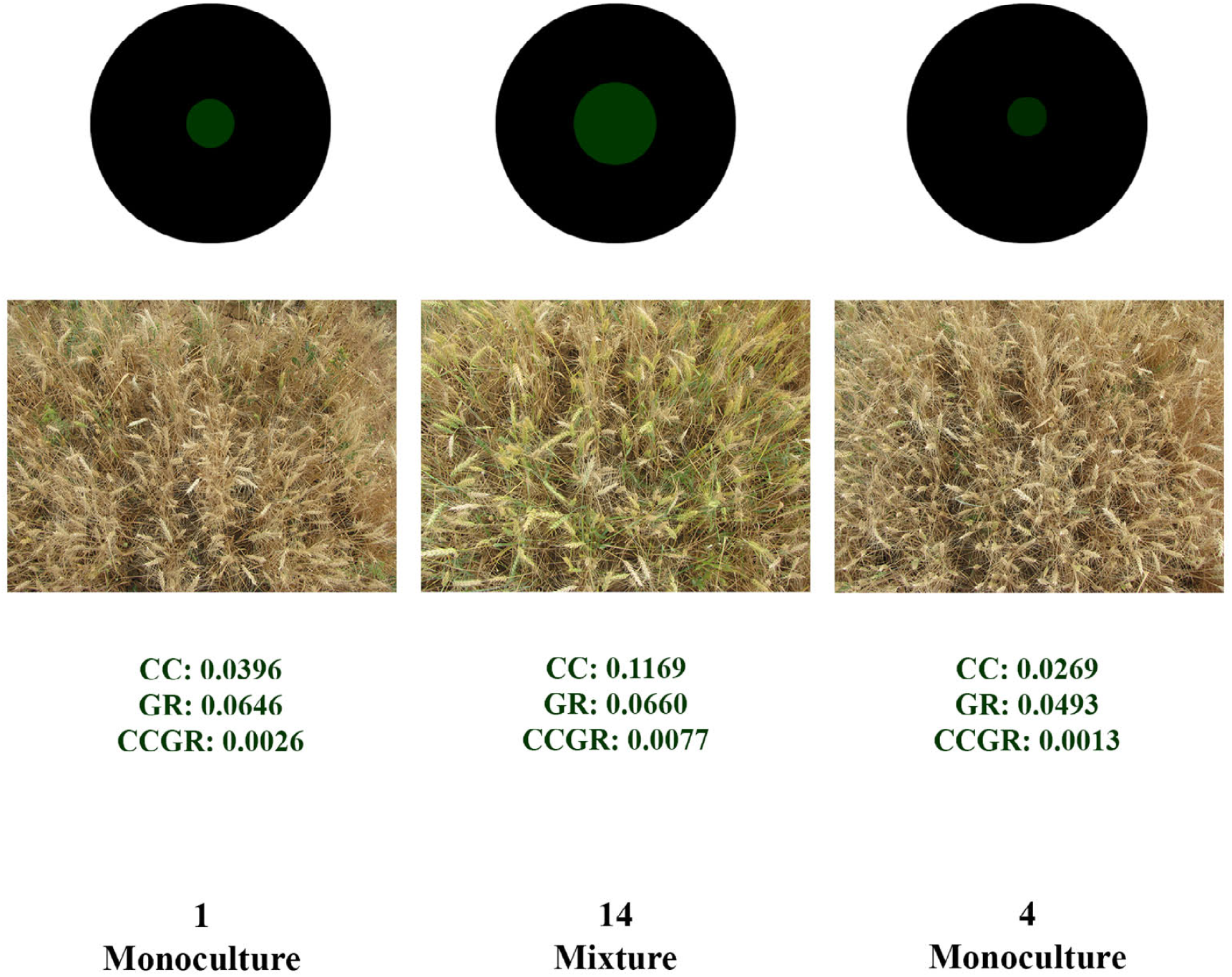
The monocultures of cultivars 1 and 4, and their mixture (14) under deficit-irrigation conditions, 197 days after sowing in the 1st year. The black circles in the background show the unit of area; and the size, and color (brightness) of the inner circles indicate CC, and GR, respectively. Notably, it is among the rare cases in which the differences in the ripening of mixture vs monocultues are easily recognisable by the original images; while in the mo & cases, it may be comparable only after estimating the image-derived criteria.

**Supplementary Figure S12.**
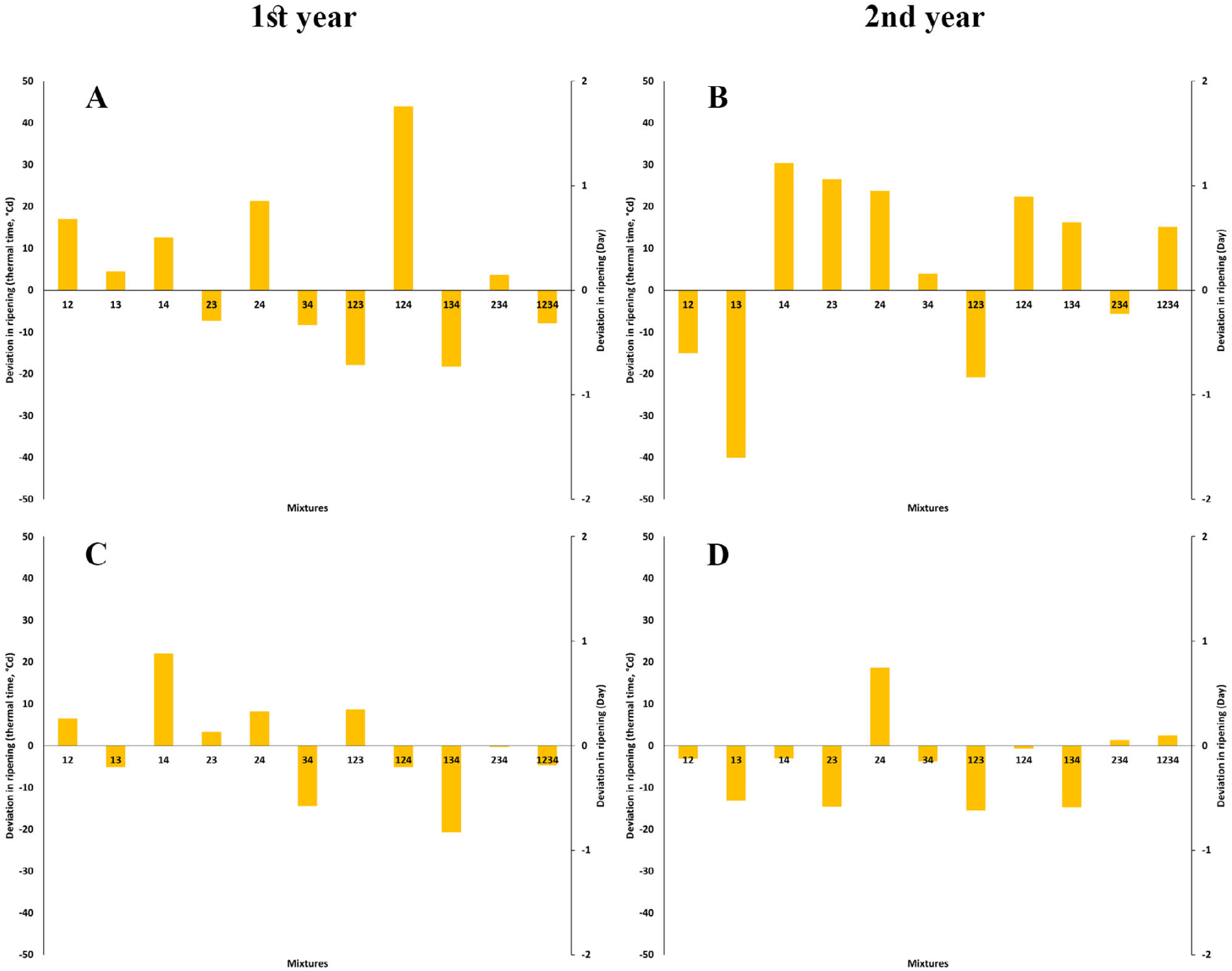
The diversion in the ripening time of mixtures from the predicted time and/or thermal time calculated based on the binomial CC trends of the respective monocultures. **A** and **B**: under well-irrigation; **C** and **D**: under deficit-irrigation. Each digit in the mixture treatments’ name show a cultivar included in the mixture.

**Supplementary S13.**
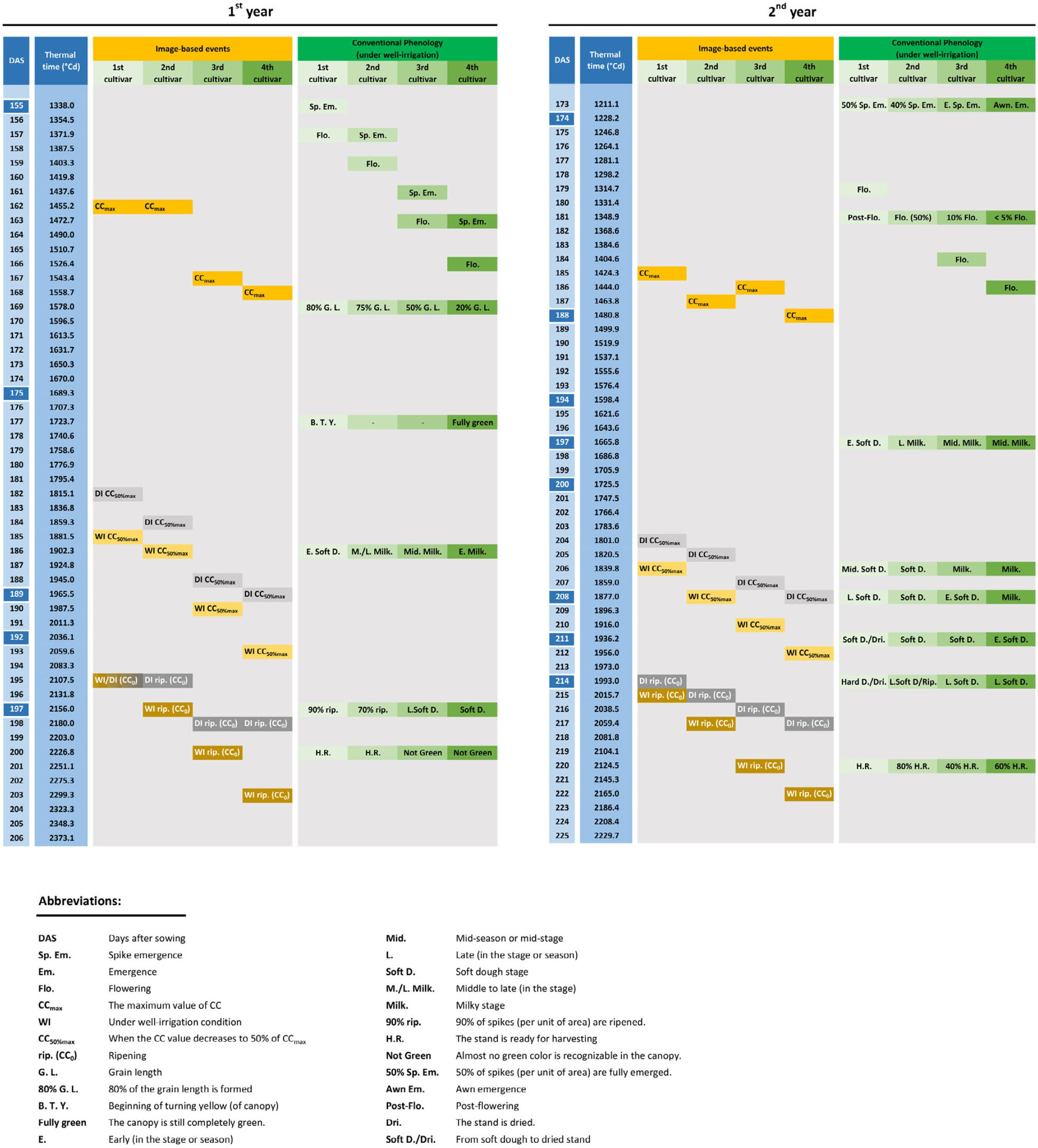
The timeline for the image-derived and conventional phenological events, during ripening of the four monocultures of wheat cultivars, over the two years of the study. The image-based events are reported using the properties of the binomial model of CC (canopy cover) declining; e.g. CC_max_ or CC_0_. The 1st to 4th cultivars are the early- to middle-ripening ones, respectively. The dark blue cells in the “DAS” column show the imaging dates.

**Supplementary Table S1.** The P-values for the effects of sources of variations (SOVs) on the image-derived indices; A and B: in the 4 monocultures during the 1st and 2nd years, respectively; C and D: in the 15 mixture treatments during the 1st and 2nd years, respectively (p<0.05). DAS: days after sowing.

| **A** |  |  |  |  |  |  |  |  |  |  |  |  |  |  |  |  |  |  |
| --- | --- | --- | --- | --- | --- | --- | --- | --- | --- | --- | --- | --- | --- | --- | --- | --- | --- | --- |
| S.O.V. | DAS= 104 |  |  | DAS= 155 |  |  | DAS= 175 |  |  | DAS= 189 |  |  | DAS= 192 |  |  | DAS= 197 |  |  |
|  | CC | GR | CCGR | CC | GR | CCGR | CC | GR | CCGR | CC | GR | CCGR | CC | GR | CCGR | CC | GR | CCGR |
| Cultivar | 0.336 | 0.000 | 0.044 | 0.360 | 0.002 | 0.056 | 0.010 | 0.006 | 0.007 | 0.000 | 0.000 | 0.000 | 0.000 | 0.000 | 0.000 | 0.000 | 0.114 | 0.001 |
| Irrigation | - | - | - | - | - | - | 0.027 | 0.003 | 0.007 | 0.000 | 0.001 | 0.000 | 0.000 | 0.001 | 0.000 | 0.007 | 0.002 | 0.005 |
| Mixtures × Irrigation | - | - | - | - | - | - | 0.101 | 0.321 | 0.258 | 0.345 | 0.132 | 0.013 | 0.029 | 0.091 | 0.003 | 0.034 | 0.086 | 0.033 |
| Replicate | 0.320 | 0.394 | 0.310 | 0.907 | 0.938 | 0.958 | 0.671 | 0.676 | 0.664 | 0.508 | 0.504 | 0.512 | 0.821 | 0.639 | 0.737 | 0.845 | 0.577 | 0.963 |

| **B** |  |  |  |  |  |  |  |  |  |  |  |  |  |  |  |  |  |  |
| --- | --- | --- | --- | --- | --- | --- | --- | --- | --- | --- | --- | --- | --- | --- | --- | --- | --- | --- |
| S.O.V. | DAS= 47 |  |  | DAS= 93 |  |  | DAS= 174 |  |  | DAS= 188 |  |  | DAS= 191 |  |  | DAS= 194 |  |  |
|  | CC | GR | CCGR | CC | GR | CCGR | CC | GR | CCGR | CC | GR | CCGR | CC | GR | CCGR | CC | GR | CCGR |
| Cultivar | 0.015 | 0.030 | 0.023 | 0.264 | 0.357 | 0.233 | 0.004 | 0.010 | 0.010 | 0.054 | 0.029 | 0.067 | 0.148 | 0.037 | 0.073 | 0.155 | 0.082 | 0.103 |
| Irrigation | - | - | - | - | - | - | - | - | - | 0.274 | 0.690 | 0.485 | 0.204 | 0.341 | 0.280 | 0.034 | 0.039 | 0.037 |
| Mixtures × Irrigation | - | - | - | - | - | - | - | - | - | 0.328 | 0.640 | 0.459 | 0.510 | 0.573 | 0.582 | 0.250 | 0.562 | 0.419 |
| Replicate | 0.462 | 0.538 | 0.569 | 0.257 | 0.877 | 0.381 | 0.991 | 0.866 | 0.965 | 0.801 | 0.147 | 0.423 | 0.563 | 0.572 | 0.566 | 0.851 | 0.750 | 0.817 |

| S.O.V. | DAS= 197 |  |  | DAS= 200 |  |  | DAS= 208 |  |  | DAS= 211 |  |  | DAS= 214 |  |  |
| --- | --- | --- | --- | --- | --- | --- | --- | --- | --- | --- | --- | --- | --- | --- | --- |
|  | CC | GR | CCGR | CC | GR | CCGR | CC | GR | CCGR | CC | GR | CCGR | CC | GR | CCGR |
| Cultivar | 0.008 | 0.001 | 0.003 | 0.006 | 0.000 | 0.002 | 0.000 | 0.003 | 0.001 | 0.000 | 0.000 | 0.000 | 0.000 | 0.003 | 0.000 |
| Irrigation | 0.001 | 0.002 | 0.001 | 0.012 | 0.027 | 0.024 | 0.000 | 0.000 | 0.000 | 0.000 | 0.000 | 0.000 | 0.000 | 0.000 | 0.000 |
| Mixtures × Irrigation | 0.265 | 0.912 | 0.744 | 0.442 | 0.621 | 0.638 | 0.843 | 0.582 | 0.612 | 0.353 | 0.446 | 0.096 | 0.009 | 0.041 | 0.002 |
| Replicate | 0.702 | 0.891 | 0.878 | 0.643 | 0.583 | 0.693 | 0.881 | 0.696 | 0.666 | 0.935 | 0.934 | 0.887 | 0.664 | 0.788 | 0.908 |

| **C** |  |  |  |  |  |  |  |  |  |  |  |  |  |  |  |  |  |  |
| --- | --- | --- | --- | --- | --- | --- | --- | --- | --- | --- | --- | --- | --- | --- | --- | --- | --- | --- |
| S.O.V. | DAS= 104 |  |  | DAS= 155 |  |  | DAS= 175 |  |  | DAS= 189 |  |  | DAS= 192 |  |  | DAS= 197 |  |  |
|  | CC | GR | CCGR | CC | GR | CCGR | CC | GR | CCGR | CC | GR | CCGR | CC | GR | CCGR | CC | GR | CCGR |
| Mixtures | 0.325 | 0.000 | 0.035 | 0.824 | 0.024 | 0.423 | 0.034 | 0.002 | 0.007 | 0.000 | 0.000 | 0.000 | 0.000 | 0.000 | 0.000 | 0.000 | 0.261 | 0.000 |
| Irrigation | - | - | - | - | - | - | 0.543 | 0.001 | 0.049 | 0.000 | 0.000 | 0.000 | 0.000 | 0.000 | 0.000 | 0.000 | 0.000 | 0.000 |
| Mixtures × Irrigation | - | - | - | - | - | - | 0.017 | 0.055 | 0.030 | 0.887 | 0.762 | 0.501 | 0.531 | 0.335 | 0.211 | 0.029 | 0.141 | 0.026 |
| Replicate | 0.033 | 0.077 | 0.027 | 0.294 | 0.473 | 0.353 | 0.064 | 0.094 | 0.056 | 0.196 | 0.959 | 0.386 | 0.022 | 0.265 | 0.032 | 0.130 | 0.422 | 0.159 |

| **D** |  |  |  |  |  |  |  |  |  |  |  |  |  |  |  |  |  |  |
| --- | --- | --- | --- | --- | --- | --- | --- | --- | --- | --- | --- | --- | --- | --- | --- | --- | --- | --- |
| S.O.V. | DAS= 47 |  |  | DAS= 93 |  |  | DAS= 174 |  |  | DAS= 188 |  |  | DAS= 191 |  |  | DAS= 194 |  |  |
|  | CC | GR | CCGR | CC | GR | CCGR | CC | GR | CCGR | CC | GR | CCGR | CC | GR | CCGR | CC | GR | CCGR |
| Mixtures | 0.000 | 0.005 | 0.000 | 0.391 | 0.313 | 0.348 | 0.001 | 0.000 | 0.003 | 0.002 | 0.000 | 0.001 | 0.000 | 0.000 | 0.000 | 0.001 | 0.002 | 0.001 |
| Irrigation | - | - | - | - | - | - | - | - | - | 0.204 | 0.954 | 0.579 | 0.114 | 0.490 | 0.321 | 0.000 | 0.000 | 0.000 |
| Mixtures × Irrigation | - | - | - | - | - | - | - | - | - | 0.560 | 0.792 | 0.721 | 0.700 | 0.747 | 0.799 | 0.379 | 0.636 | 0.570 |
| Replicate | 0.004 | 0.261 | 0.013 | 0.726 | 0.424 | 0.549 | 0.129 | 0.329 | 0.178 | 0.187 | 0.102 | 0.114 | 0.206 | 0.067 | 0.081 | 0.169 | 0.066 | 0.084 |

| S.O.V. | DAS= 197 |  |  | DAS= 200 |  |  | DAS= 208 |  |  | DAS= 211 |  |  | DAS= 214 |  |  |
| --- | --- | --- | --- | --- | --- | --- | --- | --- | --- | --- | --- | --- | --- | --- | --- |
|  | CC | GR | CCGR | CC | GR | CCGR | CC | GR | CCGR | CC | GR | CCGR | CC | GR | CCGR |
| Cultivar | 0.000 | 0.001 | 0.000 | 0.000 | 0.000 | 0.000 | 0.000 | 0.000 | 0.000 | 0.000 | 0.000 | 0.000 | 0.000 | 0.006 | 0.000 |
| Irrigation | 0.000 | 0.000 | 0.000 | 0.000 | 0.000 | 0.000 | 0.000 | 0.000 | 0.000 | 0.000 | 0.000 | 0.000 | 0.000 | 0.000 | 0.000 |
| Mixtures × Irrigation | 0.488 | 0.511 | 0.670 | 0.557 | 0.469 | 0.612 | 0.886 | 0.192 | 0.385 | 0.166 | 0.121 | 0.017 | 0.039 | 0.127 | 0.020 |
| Replicate | 0.025 | 0.009 | 0.009 | 0.042 | 0.126 | 0.050 | 0.171 | 0.085 | 0.074 | 0.288 | 0.431 | 0.288 | 0.187 | 0.208 | 0.158 |

**Supplementary Table S2.**
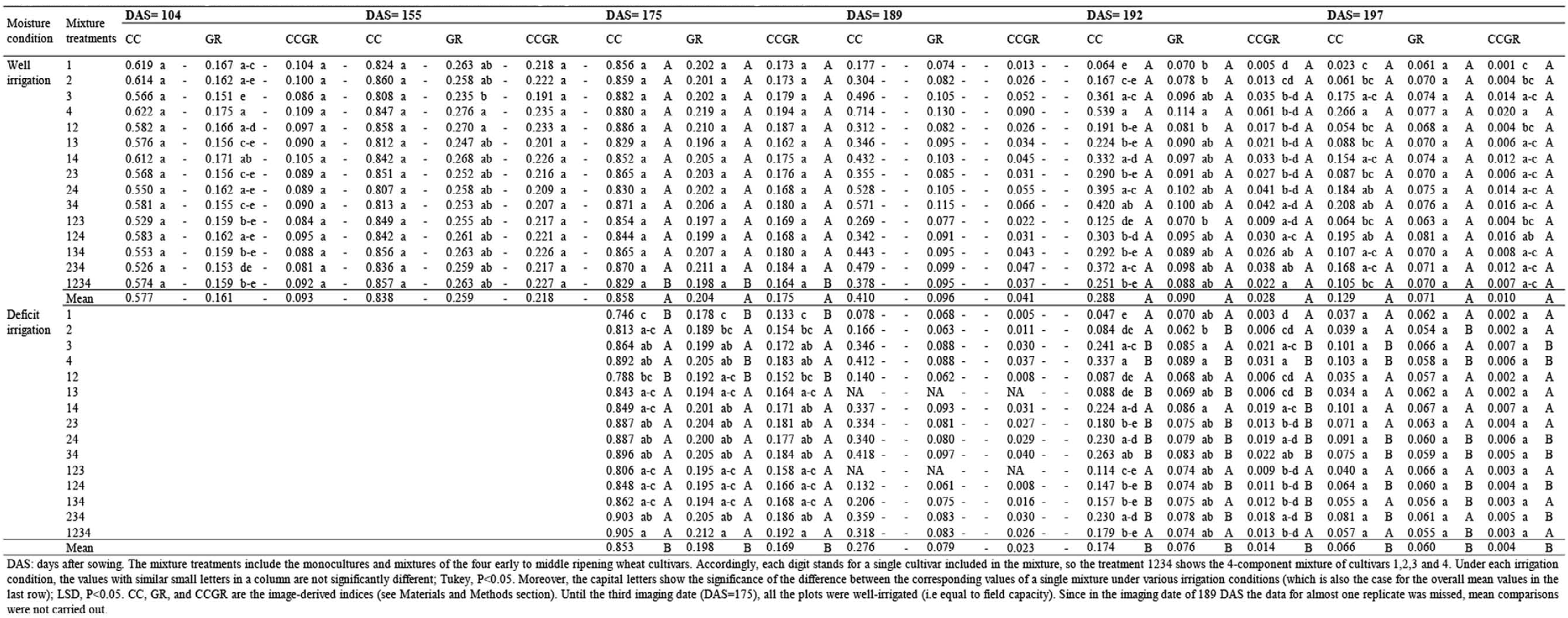
Mean comparison of the diurnal values of the image-derived indices CC, GR, and CCGR in the mixtures, during the 1st year.

**Supplementary Table S3.**
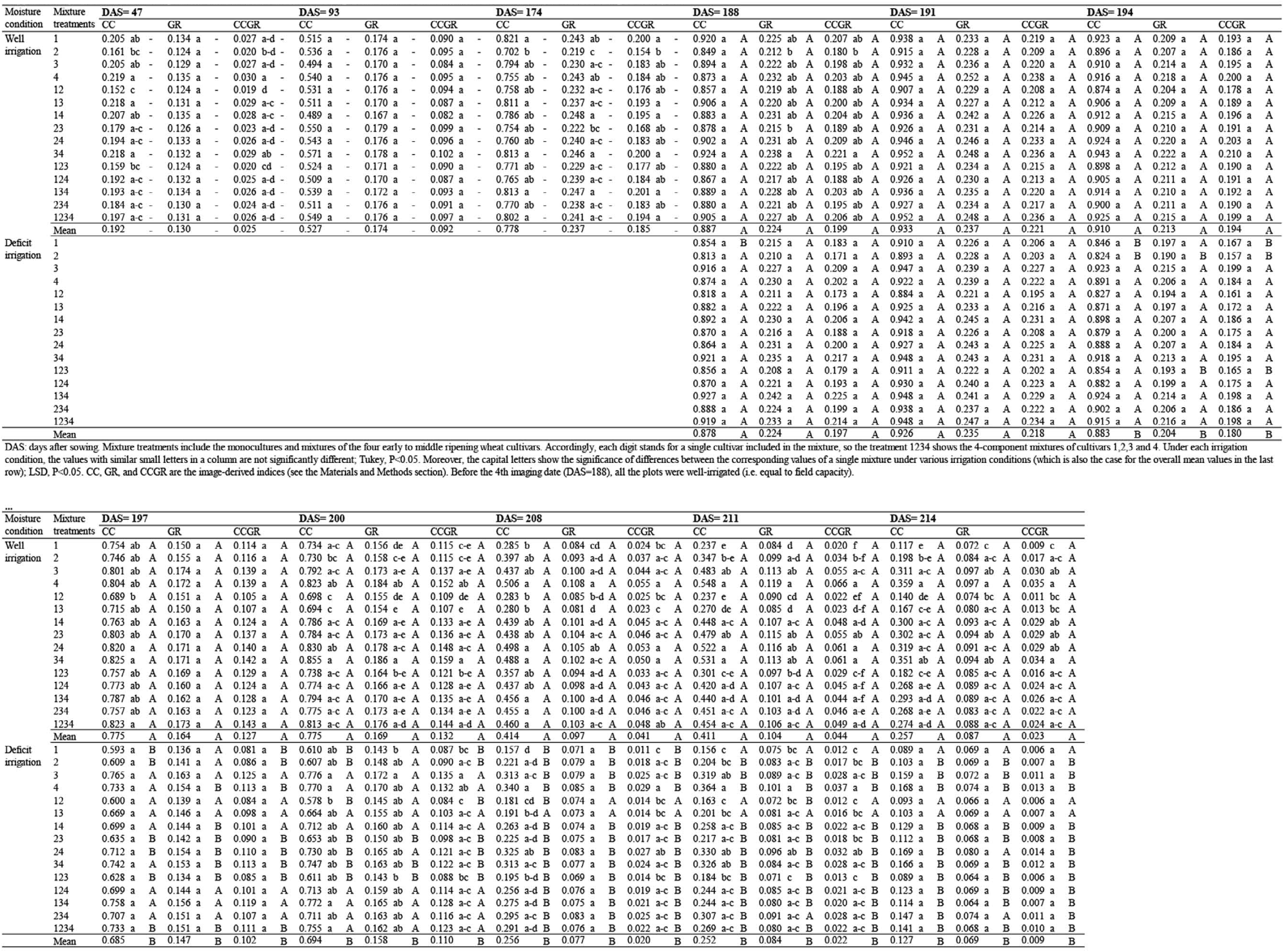
Mean comparison of the diurnal values of the image-derived iodices CC. GR. and CCGR in the mixtures, during the 2nd year.

**Supplementary Table S4.** The P-values for the effects of sources of variations (SOVs) on the binomial trends of CC against thermal time; A and B: in the 4 monocultures during the 1st and 2nd years, respectively; C and D: in the 15 mixture treatments during the 1st and 2nd years, respectively.

| **A** CC |  |  |  |  |  |  |  |  |  |
| --- | --- | --- | --- | --- | --- | --- | --- | --- | --- |
| S.O.V. | R<sup>2</sup> | RMSE | *a* | *b* | *c* | Thermal time to CC<sub>rip</sub> | Thermal time to CC<sub>max</sub> | CC<sub>max</sub> | Thermal time CC<sub>max</sub> to CC<sub>rip</sub> |
| Cultivar | **0.000** | **0.000** | 1.000 | **0.001** | **0.000** | **0.000** | **0.000** | 0.703 | **0.003** |
| Irrigation | **0.006** | **0.001** | 1.000 | **0.038** | **0.033** | **0.001** | **0.007** | 0.595 | **0.049** |
| Mixtures × Irrigation | 0.767 | 0.287 | 1.000 | 0.379 | 0.525 | **0.033** | 0.187 | 0.797 | 0.052 |
| Replicate | 0.462 | 0.495 | 1.000 | 0.788 | 0.800 | 0.988 | 0.806 | 0.832 | 0.803 |

| **B** CC |  |  |  |  |  |  |  |  |  |
| --- | --- | --- | --- | --- | --- | --- | --- | --- | --- |
| S.O.V. | R<sup>2</sup> | RMSE | *a* | *b* | *c* | Thermal time to CC<sub>rip</sub> | Thermal time to CC<sub>max</sub> | CC<sub>max</sub> | Thermal time CC<sub>max</sub> to CC<sub>rip</sub> |
| Cultivar | 0.407 | **0.046** | 1.000 | 0.286 | 0.112 | **0.000** | **0.000** | 0.057 | **0.002** |
| Irrigation | **0.037** | **0.000** | 1.000 | 0.897 | 0.588 | **0.000** | **0.000** | 0.224 | **0.016** |
| Mixtures × Irrigation | **0.002** | 0.080 | 1.000 | **0.002** | **0.009** | **0.012** | 0.949 | 0.193 | **0.004** |
| Replicate | 0.405 | 0.477 | 1.000 | 0.398 | 0.443 | 0.963 | 0.628 | 0.924 | 0.487 |

| **C** CC |  |  |  |  |  |  |  |  |  |
| --- | --- | --- | --- | --- | --- | --- | --- | --- | --- |
| S.O.V. | R<sup>2</sup> | RMSE | *a* | *b* | *c* | Thermal time to CC<sub>rip</sub> | Thermal time to CC<sub>max</sub> | CC<sub>max</sub> | Thermal time CC<sub>max</sub> to CC<sub>rip</sub> |
| Mixtures | **0.000** | **0.000** | 1.000 | **0.000** | **0.000** | **0.000** | **0.000** | 0.962 | **0.000** |
| Irrigation | **0.003** | **0.000** | 1.000 | **0.034** | **0.008** | **0.000** | **0.000** | 0.285 | **0.035** |
| Mixtures × Irrigation | 0.814 | 0.609 | 1.000 | 0.521 | 0.718 | 0.081 | 0.236 | 0.528 | **0.030** |
| Replicate | 0.109 | **0.018** | 1.000 | 0.358 | 0.401 | 0.128 | 0.245 | 0.141 | 0.426 |

| **D** CC |  |  |  |  |  |  |  |  |  |
| --- | --- | --- | --- | --- | --- | --- | --- | --- | --- |
| S.O.V. | R<sup>2</sup> | RMSE | *a* | *b* | *c* | Thermal time to CC<sub>rip</sub> | Thermal time to CC<sub>max</sub> | CC<sub>max</sub> | Thermal time CC<sub>max</sub> to CC<sub>rip</sub> |
| Mixtures | 0.387 | **0.003** | 1.000 | 0.693 | 0.373 | **0.000** | **0.000** | **0.003** | **0.008** |
| Irrigation | **0.018** | **0.000** | 1.000 | **0.013** | 0.341 | **0.000** | **0.000** | 0.130 | **0.000** |
| Mixtures × Irrigation | **0.047** | 0.365 | 1.000 | **0.001** | **0.005** | **0.018** | 0.834 | 0.331 | **0.004** |
| Replicate | 0.221 | 0.219 | 1.000 | 0.081 | 0.057 | 0.303 | 0.056 | 0.089 | 0.095 |

S.O.V: sources of variations; CC: the image-derived index of canopy cover (see the Materials and Methods section); *a*, *b*, and *c* are the equation coefficients in the general form of $CC=a(ATT)^2+b(ATT)+c$ , where the ATT is the accumulated thermal time. $Lslope_{max}$ and $Lslope_{rip}$ are the slopes of linear increasing and declining trends from sowing to the maximum amount, and then from maximum to the least values of the indices at the last imaging date (at ripening), respectively. Obviously, the ratio of $Lslope_{max}/Lslope_{rip}$ shows the comparative rate of increasing to declining trends (i.e. corresponding to the periods of reaching to the maximum values, ripening, respectively). The experimental design was RCBD; $p<0.05$ .

**Supplementary Table S5.** The properties of the binomial equations for the trends of CC (cauopy cover) against thermal time during ripening of cultivar mixtures under well and deficult irrigation conditions of the 1st year.

| Moisture condition | Mixture treatments | R<sup>2</sup> |  | RMSE |  | *a* |  | *b* |  | *c* |  | Thermal time to CC<sub>rip</sub> (°Cd) |  | Thermal time to CC<sub>max</sub> (°Cd) |  | CC<sub>max</sub> |  | Thermal time CC<sub>max</sub> to CC<sub>rip</sub> (°Cd) |  |
| --- | --- | --- | --- | --- | --- | --- | --- | --- | --- | --- | --- | --- | --- | --- | --- | --- | --- | --- | --- |
| Well irrigation | 1 | 0.905 | b A | 0.18342 | a A | -1.5E-06 | a B | 0.00404 | a A | -1.83578 | a B | 2113.26 | d A | 1345.76 | b A | 0.886 | a A | 767.49 | a B |
|  | 2 | 0.956 | ab A | 0.12076 | a-c A | -1.6E-06 | a A | 0.00462 | a A | -2.32966 | a A | 2148.25 | b-d A | 1385.56 | ab A | 0.922 | a A | 762.69 | a A |
|  | 3 | 0.983 | a A | 0.05684 | bc A | -1.9E-06 | a A | 0.00572 | a A | -3.45345 | a A | 2216.06 | a-c A | 1518.57 | ab A | 0.899 | a A | 697.49 | a A |
|  | 4 | 0.995 | a A | 0.02866 | c A | -1.9E-06 | a A | 0.00581 | a A | -3.64481 | a A | 2273.26 | a A | 1568.95 | a A | 0.918 | a A | 704.31 | a A |
|  | 12 | 0.961 | ab A | 0.10668 | a-c A | -1.9E-06 | a B | 0.00542 | a A | -3.00377 | a B | 2147.51 | b-d A | 1440.24 | ab A | 0.927 | a A | 707.27 | a B |
|  | 13 | 0.950 | ab A | 0.09997 | a-c B | -1.8E-06 | a A | 0.00538 | a A | -3.18529 | a A | 2164.27 | b-d A | 1480.60 | ab A | 0.829 | a A | 683.67 | a A |
|  | 14 | 0.962 | ab A | 0.08104 | a-c A | -1.7E-06 | a A | 0.00501 | a A | -2.84811 | a A | 2204.30 | a-c A | 1473.55 | ab A | 0.879 | a A | 730.75 | a A |
|  | 23 | 0.979 | ab A | 0.05987 | bc B | -2.0E-06 | a A | 0.00598 | a A | -3.60033 | a A | 2174.83 | b-d A | 1504.57 | ab A | 0.896 | a A | 670.25 | a A |
|  | 24 | 0.996 | a A | 0.02382 | c B | -1.9E-06 | a A | 0.00585 | a A | -3.69181 | a A | 2226.51 | a-c A | 1553.69 | ab A | 0.852 | a A | 672.82 | a A |
|  | 34 | 0.995 | a A | 0.02443 | c B | -2.0E-06 | a A | 0.00623 | a A | -3.98608 | a A | 2232.93 | ab A | 1564.28 | a A | 0.890 | a A | 668.65 | a A |
|  | 123 | 0.928 | ab A | 0.14848 | ab A | -1.5E-06 | a A | 0.00416 | a A | -1.96770 | a A | 2138.65 | cd A | 1357.70 | ab A | 0.902 | a A | 780.95 | a A |
|  | 124 | 0.955 | ab A | 0.09162 | a-c B | -1.4E-06 | a A | 0.00420 | a A | -2.18418 | a A | 2221.71 | a-c A | 1441.06 | ab A | 0.870 | a A | 780.65 | a A |
|  | 134 | 0.969 | ab A | 0.06520 | bc B | -1.8E-06 | a A | 0.00526 | a A | -2.95780 | a A | 2184.00 | a-d A | 1446.73 | ab A | 0.919 | a A | 737.27 | a A |
|  | 234 | 0.984 | a A | 0.05198 | bc B | -1.9E-06 | a A | 0.00590 | a A | -3.63679 | a A | 2214.28 | a-c A | 1529.29 | ab A | 0.893 | a A | 684.99 | a A |
|  | 1234 | 0.965 | ab A | 0.08968 | a-c A | -1.7E-06 | a A | 0.00514 | a A | -2.95452 | a A | 2176.10 | b-d A | 1471.03 | ab A | 0.850 | a B | 705.07 | a A |
|  | Mean | 0.966 | A | 0.08216 | B | -1.8E-06 | A | 0.00525 | A | -3.01867 | B | 2189.06 | A | 1472.11 | A | 0.889 | A | 716.96 |  |
| Deficit irrigation | 1 | 0.884 | b A | 0.19049 | a A | -8.8E-07 | a A | 0.00195 | c B | -0.18906 | a A | 2109.24 | c A | 1091.73 | b B | 0.903 | a A | 1017.51 | a A |
|  | 2 | 0.923 | ab A | 0.17082 | ab A | -1.4E-06 | ab A | 0.00373 | bc A | -1.62710 | ab A | 2125.42 | bc A | 1316.86 | ab A | 0.884 | a A | 808.56 | b A |
|  | 3 | 0.947 | ab A | 0.10860 | a-c A | -1.8E-06 | bc A | 0.00540 | ab A | -3.14973 | bc A | 2167.69 | ab B | 1479.88 | a A | 0.855 | a A | 687.81 | b A |
|  | 4 | 0.981 | ab A | 0.06681 | c A | -1.9E-06 | bc A | 0.00548 | ab A | -3.12934 | bc A | 2182.73 | a B | 1477.08 | a A | 0.924 | a A | 705.64 | b A |
|  | 12 | 0.931 | ab A | 0.15790 | ab A | -1.3E-06 | ab A | 0.00333 | bc B | -1.30243 | ab A | 2126.75 | bc A | 1292.43 | ab B | 0.878 | a A | 834.32 | ab A |
|  | 13 | 0.946 | ab A | 0.18396 | a A | -1.8E-06 | bc A | 0.00501 | ab A | -2.66513 | bc A | 2134.35 | a-c A | 1421.13 | a A | 0.898 | a A | 713.22 | b A |
|  | 14 | 0.962 | ab A | 0.11632 | a-c A | -1.6E-06 | bc A | 0.00469 | ab A | -2.46300 | bc A | 2172.73 | ab A | 1432.33 | a A | 0.895 | a A | 740.40 | b A |
|  | 23 | 0.953 | ab A | 0.13227 | a-c A | -1.8E-06 | bc A | 0.00522 | ab A | -2.84983 | bc A | 2151.43 | a-c A | 1440.65 | a A | 0.914 | a A | 710.79 | b A |
|  | 24 | 0.956 | ab A | 0.11393 | a-c A | -1.9E-06 | bc A | 0.00543 | ab A | -3.07220 | bc A | 2163.26 | ab B | 1458.86 | a A | 0.909 | a A | 704.40 | b A |
|  | 34 | 0.980 | ab A | 0.08996 | bc A | -2.1E-06 | c A | 0.00649 | a A | -3.98913 | c A | 2165.38 | ab B | 1511.50 | a A | 0.919 | a A | 653.88 | b A |
|  | 123 | 0.957 | a A | 0.15496 | a-c A | -1.7E-06 | bc A | 0.00468 | ab A | -2.44670 | bc A | 2141.04 | a-c A | 1411.93 | a A | 0.870 | a A | 729.12 | b A |
|  | 124 | 0.944 | a A | 0.15342 | a-c A | -1.6E-06 | bc A | 0.00450 | ab A | -2.25745 | a-c A | 2147.03 | a-c B | 1392.61 | a A | 0.902 | a A | 754.42 | b A |
|  | 134 | 0.943 | a A | 0.15210 | a-c A | -1.8E-06 | bc A | 0.00513 | ab A | -2.80467 | bc A | 2144.03 | a-c A | 1429.73 | a A | 0.895 | a A | 714.30 | b A |
|  | 234 | 0.961 | a A | 0.11038 | a-c A | -1.9E-06 | bc A | 0.00572 | ab A | -3.29194 | bc A | 2160.22 | a-c B | 1467.32 | a A | 0.922 | a A | 692.90 | b A |
|  | 1234 | 0.960 | a A | 0.13787 | a-c A | -1.9E-06 | bc A | 0.00530 | ab A | -2.83627 | bc A | 2148.19 | a-c A | 1427.99 | a A | 0.957 | a A | 720.20 | b A |
|  | Mean | 0.949 | B | 0.13599 | A | -1.7E-06 | A | 0.00481 | B | -2.53827 | A | 2149.30 | B | 1403.47 | B | 0.902 | A | 745.83 |  |

Mixture treatments include the monocultures and mixtures of the four early to middle ripening wheat cultivars. Accordingly, each digit stands for a single cultivar included in the mixture, so the treatment 1234 shows the 4-component mixtures of cultivars 1,2,3 and 4. The R<sup>2</sup> coefficient of determination, and RMSE shows the model goodness of fitness; *a*, *b*, and *c* are the coefficients of equation for the binomial model (in the general form of $CC=a(ATT)^2+b(ATT)+c$ , where the ATT is the accumulated thermal time). CC<sub>rip</sub> and CC<sub>max</sub> are the model-based estimated amounts of CC equal to zero (at ripening), and the maximum value at the peak, respectively. Under each irrigation condition, the values with similar small letters in a column are not significantly different; Tukey, P<0.05. Moreover, the capital letters show the significance of differences between the corresponding values of a single mixture under various irrigation conditions (which is also the case for the overall mean values in the last row); LSD, P<0.05.

**Supplementary Table S6.** The properties of binomial equations for the trends of CC (canopy cover) against thermal time during ripening of cultivator mixtures under well- and deficit-irrigation conditions of the 2nd year.

| Moisture condition | Mixture treatments | R<sup>2</sup> |  | RMSE |  | *a* |  | *b* |  | *c* |  | Thermal time to CC<sub>rip</sub> (°Cd) |  | Thermal time to CC<sub>max</sub> (°Cd) |  | CC<sub>max</sub> |  | Thermal time CC<sub>max</sub> to CC<sub>rip</sub> (°Cd) |  |
| --- | --- | --- | --- | --- | --- | --- | --- | --- | --- | --- | --- | --- | --- | --- | --- | --- | --- | --- | --- |
| Well irrigation | 1 | 0.976 | a A | 0.05915 | a B | -2.7E-06 | b B | 0.00776 | a A | -4.56575 | a B | 2013.58 | f A | 1422.62 | c A | 0.952 | a A | 590.96 | bc A |
|  | 2 | 0.976 | a A | 0.04692 | a B | -2.7E-06 | ab A | 0.00788 | a A | -4.94107 | a A | 2053.78 | b-f A | 1477.35 | ab A | 0.884 | a A | 576.43 | c A |
|  | 3 | 0.955 | a A | 0.05653 | a A | -2.3E-06 | ab A | 0.00688 | a A | -4.17612 | a A | 2104.48 | a-c A | 1478.43 | ab A | 0.911 | a A | 626.04 | a-c A |
|  | 4 | 0.944 | a A | 0.05559 | a A | -2.2E-06 | a A | 0.00652 | a B | -3.96091 | a A | 2138.92 | a A | 1493.54 | a A | 0.910 | a A | 645.38 | ab A |
|  | 12 | 0.964 | a A | 0.06569 | a B | -2.5E-06 | ab A | 0.00728 | a A | -4.31839 | a A | 2021.38 | ef A | 1429.36 | bc A | 0.890 | a A | 592.02 | bc A |
|  | 13 | 0.959 | a A | 0.07157 | a A | -2.5E-06 | ab A | 0.00709 | a A | -4.11352 | a A | 2027.26 | d-f A | 1419.43 | c A | 0.922 | a A | 607.83 | a-c A |
|  | 14 | 0.957 | a A | 0.05500 | a B | -2.3E-06 | ab A | 0.00666 | a A | -3.97684 | a A | 2102.02 | a-d A | 1468.17 | a-c A | 0.908 | a A | 633.85 | a-c A |
|  | 23 | 0.958 | a A | 0.05442 | a B | -2.4E-06 | ab A | 0.00717 | a A | -4.43907 | a A | 2102.25 | a-c A | 1489.23 | a A | 0.902 | a A | 613.02 | a-c A |
|  | 24 | 0.954 | a A | 0.05298 | a A | -2.3E-06 | ab A | 0.00678 | a B | -4.08976 | a A | 2119.28 | ab A | 1481.01 | ab A | 0.931 | a A | 638.27 | a-c A |
|  | 34 | 0.946 | a A | 0.05808 | a A | -2.2E-06 | a A | 0.00644 | a B | -3.79085 | a A | 2130.03 | a A | 1471.24 | a-c A | 0.948 | a A | 658.79 | a A |
|  | 123 | 0.976 | a A | 0.04983 | a B | -2.6E-06 | ab A | 0.00766 | a A | -4.65868 | a A | 2041.49 | c-f A | 1452.06 | a-c A | 0.910 | a A | 589.43 | bc A |
|  | 124 | 0.971 | a A | 0.04758 | a B | -2.4E-06 | ab A | 0.00703 | a A | -4.27323 | a A | 2086.11 | a-f A | 1471.52 | a-c A | 0.903 | a A | 614.59 | a-c A |
|  | 134 | 0.970 | a A | 0.04619 | a B | -2.3E-06 | ab A | 0.00670 | a B | -3.98945 | a A | 2098.01 | a-d A | 1464.20 | a-c A | 0.919 | a A | 633.81 | a-c A |
|  | 234 | 0.967 | a A | 0.04842 | a B | -2.4E-06 | ab A | 0.00700 | a A | -4.27636 | a A | 2093.34 | a-e A | 1476.11 | ab A | 0.900 | a A | 617.23 | a-c A |
|  | 1234 | 0.971 | a A | 0.04791 | a B | -2.5E-06 | ab A | 0.00741 | a A | -4.54091 | a A | 2088.94 | a-e A | 1477.66 | ab A | 0.936 | a A | 611.28 | a-c A |
|  | Mean | 0.963 | A | 0.05439 | B | -2.42E-06 | A | 0.00709 | B | -4.27406 | A | 2081.39 | A | 1464.80 | A | 0.915 | A | 616.59 |  |
| Deficit irrigation | 1 | 0.941 | a B | 0.09416 | a A | -2.3E-06 | a A | 0.00647 | b B | -3.57603 | a A | 1991.44 | c A | 1377.63 | b B | 0.884 | a B | 613.82 | a A |
|  | 2 | 0.948 | a B | 0.07906 | a A | -2.6E-06 | ab A | 0.00736 | ab A | -4.45587 | ab A | 2004.86 | a-c B | 1435.10 | ab B | 0.830 | a A | 569.75 | a A |
|  | 3 | 0.964 | a A | 0.06713 | a A | -2.6E-06 | ab B | 0.00761 | ab A | -4.50880 | ab A | 2033.05 | a-c B | 1433.54 | ab B | 0.950 | a A | 599.50 | a A |
|  | 4 | 0.958 | a A | 0.06619 | a A | -2.6E-06 | ab B | 0.00771 | ab A | -4.73420 | ab B | 2045.42 | a B | 1461.17 | a A | 0.902 | a A | 584.25 | a B |
|  | 12 | 0.945 | a A | 0.08513 | a A | -2.4E-06 | ab A | 0.00670 | ab A | -3.85594 | ab A | 1996.10 | bc A | 1401.79 | ab A | 0.843 | a A | 594.31 | a A |
|  | 13 | 0.953 | a A | 0.08356 | a A | -2.5E-06 | ab A | 0.00705 | ab A | -4.02650 | ab A | 2000.87 | a-c A | 1398.98 | ab A | 0.911 | a A | 601.89 | a A |
|  | 14 | 0.959 | a A | 0.07250 | a A | -2.6E-06 | ab B | 0.00750 | ab A | -4.43296 | ab A | 2017.30 | a-c B | 1426.31 | ab B | 0.919 | a A | 590.99 | a B |
|  | 23 | 0.947 | a A | 0.08371 | a A | -2.5E-06 | ab A | 0.00715 | ab A | -4.16920 | ab A | 2007.93 | a-c B | 1414.71 | ab B | 0.887 | a A | 593.23 | a A |
|  | 24 | 0.957 | a A | 0.06623 | a A | -2.7E-06 | ab B | 0.00774 | ab A | -4.76723 | ab A | 2039.54 | ab B | 1460.42 | a A | 0.888 | a A | 579.12 | a B |
|  | 34 | 0.962 | a A | 0.06704 | a A | -2.6E-06 | ab B | 0.00752 | ab A | -4.46772 | ab A | 2035.85 | a-c B | 1437.39 | ab A | 0.936 | a A | 598.46 | a B |
|  | 123 | 0.951 | a B | 0.08279 | a A | -2.5E-06 | ab A | 0.00700 | ab A | -4.04042 | ab A | 1996.33 | bc B | 1400.55 | ab B | 0.880 | a A | 595.78 | a A |
|  | 124 | 0.959 | a A | 0.07240 | a A | -2.7E-06 | ab B | 0.00776 | ab A | -4.67433 | ab A | 2014.25 | a-c B | 1436.46 | ab B | 0.902 | a A | 577.79 | a B |
|  | 134 | 0.970 | a A | 0.06677 | a A | -2.8E-06 | b B | 0.00812 | a A | -4.84646 | b B | 2012.22 | a-c B | 1430.09 | ab A | 0.959 | a A | 582.13 | a B |
|  | 234 | 0.961 | a A | 0.06831 | a A | -2.5E-06 | ab A | 0.00730 | ab A | -4.31243 | ab A | 2030.52 | a-c B | 1431.62 | ab B | 0.913 | a A | 598.90 | a A |
|  | 1234 | 0.966 | a A | 0.06752 | a A | -2.7E-06 | ab A | 0.00767 | ab A | -4.53663 | ab A | 2022.15 | a-c B | 1428.04 | ab B | 0.945 | a A | 594.11 | a A |
|  | Mean | 0.956 | B | 0.07483 | A | -2.59E-06 | A | 0.00738 | A | -4.36031 | A | 2016.52 | B | 1424.92 | B | 0.903 | A | 591.60 |  |

Mixture treatments include the monocultures and mixtures of the four early to middle ripening wheat cultivars. Accordingly, each digit stands for a single cultivar included in the mixture, so the treatment 1234 shows the 4-component mixtures of cultivars 1,2,3 and 4. The R<sup>2</sup> coefficient of determination, and RMSE shows the model goodness of fitness; *a*, *b*, and *c* are the coefficients of equation for the binomial model (in the general form of $CC = a(ATT)^2 + b(ATT) + c$ , where the ATT is the accumulated thermal time). CC<sub>rip</sub> and CC<sub>max</sub> are the model-based estimated amounts of CC equal to zero (at ripening), and the maximum value at the peak, respectively. Under each irrigation condition, the values with similar small letters in a column are not significantly different; Tukey, P<0.05. Moreover, the capital letters show the significance of differences between the corresponding values of a single mixture under various irrigation conditions (which is also the case for the overall mean values in the last row); LSD, P<0.05.

**Supplementary Table S7.** The P-values for the effects of sources of variations (SOVs) on the properties of the linear trends of image-derived indices (CC, GR, and CCGR) against thermal time. A and B: in the 4 mono cultures during the 1st and 2nd years, respectively; C and D: in the 15 mixture treatments during the 1st and 2nd yean, respectively.

| S.O.V. | CC |  |  | GR |  |  | CCGR |  |  |
| --- | --- | --- | --- | --- | --- | --- | --- | --- | --- |
|  | LSlope<sub>max</sub> | LSlope<sub>rip.</sub> | Lslope<sub>rip.</sub> /LSlope<sub>max.</sub> | LSlope<sub>max</sub> | LSlope<sub>rip.</sub> | Lslope<sub>rip.</sub> /LSlope<sub>max.</sub> | LSlope<sub>max</sub> | LSlope<sub>rip.</sub> | Lslope<sub>rip.</sub> /LSlope<sub>max.</sub> |
| Cultivar | **0.010** | 0.114 | **0.000** | **0.006** | 0.140 | 0.467 | **0.007** | 0.100 | **0.001** |
| Irrigation | **0.027** | 0.406 | **0.008** | **0.003** | 0.544 | 0.087 | **0.007** | **0.044** | **0.010** |
| Mixtures × Irrigation | 0.101 | **0.005** | **0.025** | 1.000 | 0.110 | 0.079 | 0.258 | 0.084 | **0.042** |
| Replicate | 0.671 | 0.984 | 0.924 | 1.000 | 0.540 | 0.528 | 1.000 | 0.711 | 0.967 |

**B**
| S.O.V. | CC |  |  | GR |  |  | CCGR |  |  |
| --- | --- | --- | --- | --- | --- | --- | --- | --- | --- |
|  | LSlope<sub>max</sub> | LSlope<sub>rip.</sub> | Lslope<sub>rip.</sub> /LSlope<sub>max.</sub> | LSlope<sub>max</sub> | LSlope<sub>rip.</sub> | Lslope<sub>rip.</sub> /LSlope<sub>max.</sub> | LSlope<sub>max</sub> | LSlope<sub>rip.</sub> | Lslope<sub>rip.</sub> /LSlope<sub>max.</sub> |
| Cultivar | 0.148 | **0.000** | **0.000** | **0.037** | 0.266 | **0.016** | 0.073 | 0.396 | **0.000** |
| Irrigation | 0.204 | **0.000** | **0.000** | 0.341 | **0.005** | **0.000** | 0.280 | 0.260 | **0.000** |
| Mixtures × Irrigation | 0.510 | **0.000** | **0.004** | 1.000 | 0.060 | **0.016** | 0.582 | 0.160 | **0.000** |
| Replicate | 0.563 | 0.978 | 0.598 | 1.000 | 0.759 | 0.913 | 0.566 | 0.522 | 0.794 |

**C**
| S.O.V. | CC |  |  | GR |  |  | CCGR |  |  |
| --- | --- | --- | --- | --- | --- | --- | --- | --- | --- |
|  | LSlope<sub>max</sub> | LSlope<sub>rip.</sub> | Lslope<sub>rip.</sub> /LSlope<sub>max.</sub> | LSlope<sub>max</sub> | LSlope<sub>rip.</sub> | Lslope<sub>rip.</sub> /LSlope<sub>max.</sub> | LSlope<sub>max</sub> | LSlope<sub>rip.</sub> | Lslope<sub>rip.</sub> /LSlope<sub>max.</sub> |
| Mixtures | **0.034** | 0.122 | **0.000** | 1.000 | 0.126 | 0.482 | **0.007** | 0.176 | **0.000** |
| Irrigation | 0.543 | **0.000** | **0.000** | **0.001** | **0.042** | **0.000** | **0.049** | 1.000 | **0.000** |
| Mixtures × Irrigation | **0.017** | **0.001** | **0.021** | 1.000 | **0.047** | 0.095 | **0.030** | **0.011** | **0.034** |
| Replicate | 0.064 | 0.915 | 0.219 | 1.000 | 0.499 | 0.718 | 0.056 | 0.169 | 0.336 |

**D**
| S.O.V. | CC |  |  | GR |  |  | CCGR |  |  |
| --- | --- | --- | --- | --- | --- | --- | --- | --- | --- |
|  | LSlope<sub>max</sub> | LSlope<sub>rip.</sub> | Lslope<sub>rip.</sub> /LSlope<sub>max.</sub> | LSlope<sub>max</sub> | LSlope<sub>rip.</sub> | Lslope<sub>rip.</sub> /LSlope<sub>max.</sub> | LSlope<sub>max</sub> | LSlope<sub>rip.</sub> | Lslope<sub>rip.</sub> /LSlope<sub>max.</sub> |
| Mixtures | **0.000** | **0.000** | **0.000** | **0.000** | 0.138 | 0.203 | **0.000** | 0.090 | **0.000** |
| Irrigation | 0.114 | **0.000** | **0.000** | 1.000 | **0.000** | **0.000** | 1.000 | **0.000** | **0.000** |
| Mixtures × Irrigation | 0.700 | **0.003** | **0.030** | 1.000 | 0.224 | 0.102 | 1.000 | 0.268 | **0.022** |
| Replicate | 0.206 | 0.531 | 0.240 | 0.067 | 0.515 | 0.623 | 0.081 | 0.316 | 0.319 |

S.O.V: sources of variations; CC, GR, and CCGR are the image-derived indices (see the Materials and Methods section); LSlope<sub>max</sub> and LSlope<sub>rip.</sub> are the slopes of the linear increasing and declining trends from sowing to the maximum observed amount, and then from the maximum to the least values of the indices at the last imaging date (at ripening), respectively. Obviously, the ratio of Lslope<sub>rip.</sub> /LSlope<sub>max.</sub> shows the comparative rate of increasing to declining trends.

**Supplementary Table S8.**
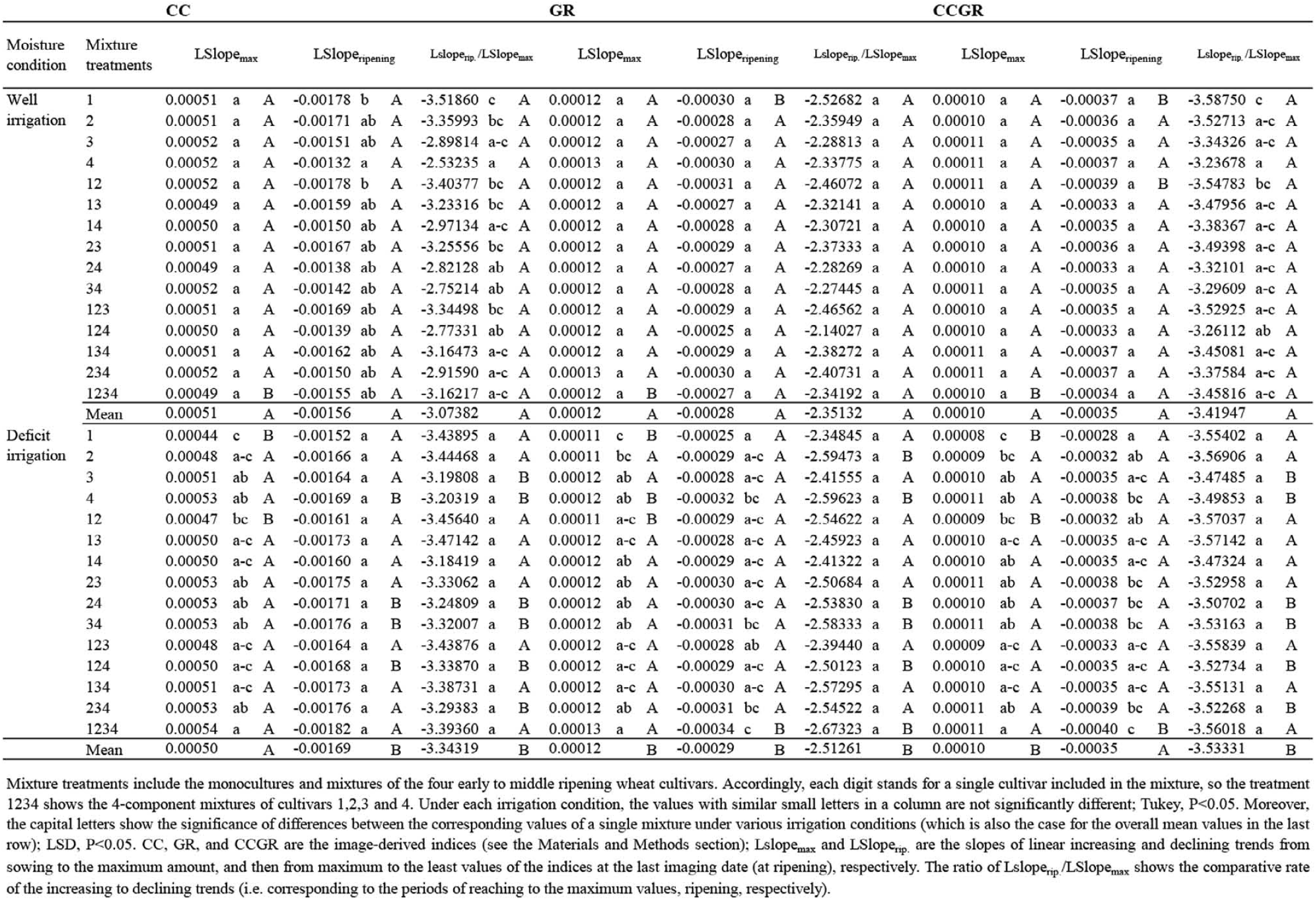
Mean comparison of the properties of the linear increasing and declining trends of the image-derived indices (CC, GR, and CCGR) in the mixtures, during the 1st year.

**Supplementary Table S9.**
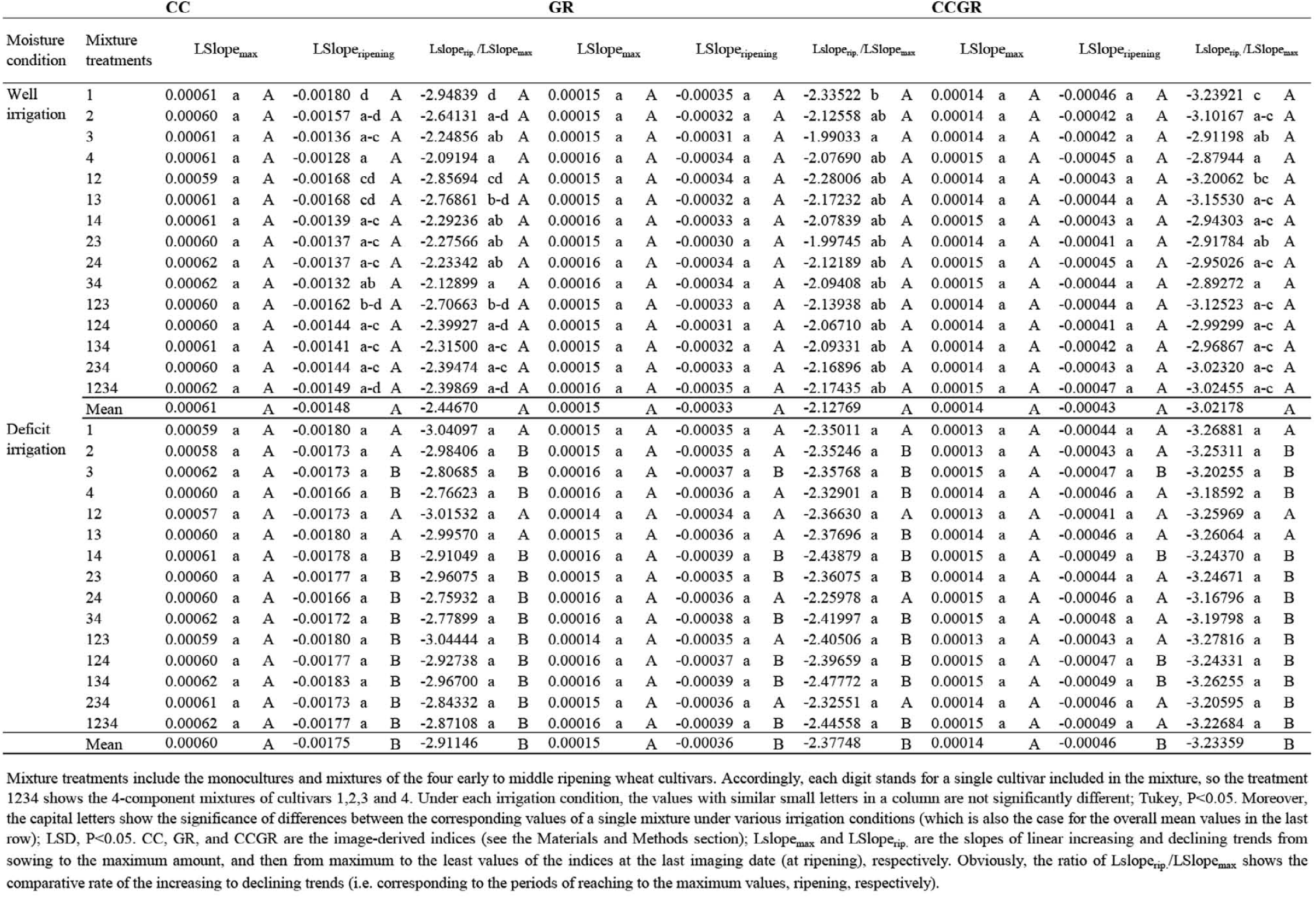
Mean comparison of the properties of the linear increasing and declining trends of the image-derived indices (CC, GR, and CCGR) in the mixtures, during the 2nd year.

**Supplementary Table S10.** The parameters of relationship between CC and GR.

| Year | DAS | Well-irrigation |  |  |  | Deficit-irrigation |  |  |  |
| --- | --- | --- | --- | --- | --- | --- | --- | --- | --- |
|  |  | Correlation |  | Linear regression |  | Correlation |  | Linear regression |  |
|  |  | R | P-value | R<sup>2</sup> | RMSE | R | P-value | R<sup>2</sup> | RMSE |
| 1st year | 104 | 0.708 | 0.003166 | 0.501 | 0.0048 | - | - | - | - |
|  | 155 | 0.581 | 0.02318 | 0.337 | 0.0083 | - | - | - | - |
|  | 175 | 0.707 | 0.003201 | 0.500 | 0.0046 | 0.918 | 1.38E-06 | 0.843 | 0.0034 |
|  | 189 | 0.970 | 2.4E-09 | 0.941 | 0.0038 | 0.908 | 2.87E-06 | 0.825 | 0.0052 |
|  | 192 | 0.977 | 4.43E-10 | 0.954 | 0.0027 | 0.910 | 2.5E-06 | 0.828 | 0.0032 |
|  | 197 | 0.874 | 2.09E-05 | 0.763 | 0.0027 | 0.318 | 0.248385 | 0.101 | 0.0040 |
| 2nd year | 47 | 0.837 | 0.000101 | 0.700 | 0.0022 | - | - | - | - |
|  | 93 | 0.796 | 0.000382 | 0.634 | 0.0022 | - | - | - | - |
|  | 174 | 0.686 | 0.004722 | 0.471 | 0.0066 | - | - | - | - |
|  | 188 | 0.630 | 0.011752 | 0.397 | 0.0058 | 0.848 | 6.41E-05 | 0.720 | 0.0056 |
|  | 191 | 0.807 | 0.000273 | 0.652 | 0.0050 | 0.859 | 4.18E-05 | 0.737 | 0.0046 |
|  | 194 | 0.816 | 0.000205 | 0.666 | 0.0029 | 0.938 | 2.44E-07 | 0.879 | 0.0031 |
|  | 197 | 0.861 | 3.82E-05 | 0.741 | 0.0046 | 0.909 | 2.73E-06 | 0.826 | 0.0035 |
|  | 200 | 0.963 | 8.4E-09 | 0.928 | 0.0028 | 0.960 | 1.44E-08 | 0.922 | 0.0028 |
|  | 208 | 0.973 | 1.28E-09 | 0.946 | 0.0020 | 0.782 | 0.000571 | 0.612 | 0.0030 |
|  | 211 | 0.949 | 6.52E-08 | 0.901 | 0.0037 | 0.883 | 1.26E-05 | 0.781 | 0.0040 |
|  | 214 | 0.930 | 5.14E-07 | 0.865 | 0.0029 | 0.715 | 0.002761 | 0.511 | 0.0030 |

DAS: days after sowing.

